# Hyperglycaemia reversibly disrupts circadian behaviour in mice

**DOI:** 10.64898/2026.10.07.757333

**Authors:** Matthew Lloyd, Guohao Ni, Kai-Ting Hu, Russell G Foster, Frances M Ashcroft, Aarti Jagannath, Sridhar R Vasudevan

**Affiliations:** Department of Pharmacology, University of Oxford, Mansfield Road, Oxford, OX1 3QT; Nuffield Department of Clinical Neurosciences, University of Oxford, New Biochemistry Building, South Parks Road, Oxford, OX1 3QU; Department of Physiology, Anatomy and Genetics, University of Oxford, Parks Road, Oxford, OX1 3PT

## Abstract

Disrupted sleep and circadian rhythms are among the most common features of diabetes, and are known to raise the risk of developing the disease. Whether the relationship also runs the other way, with diabetes itself disrupting the clock, has remained unresolved. Using an inducible mouse model of diabetes, in which hyperglycaemia can be induced and then reversed within the same animal, we show that elevated blood glucose is a reversible cause of circadian disruption. Raising blood glucose reduced the amplitude of circadian rest-activity rhythms by 50%, in proportion to the degree of hyperglycaemia, and shortened the free-running period of the clock. Restoring normoglycaemia reversed both effects. Diabetic animals had a delayed phase of activity onset, i.e. later chronotype, and re-entrained more slowly after a simulated jet lag. However, phase shifting to nocturnal light pulses was intact, suggesting normal photic responses. Brain glucose and O-GlcNAcylation, a nutrient-sensitive protein modification, were both elevated in diabetic animals, showing that peripheral hyperglycaemia reaches the brain and alters its metabolic state, and providing a molecular readout of that exposure. Together, these findings establish that a peripheral metabolic state is a reversible determinant of circadian rest-activity behaviour, completing a bidirectional link between blood glucose and the clock and arguing that the two must be treated together rather than in isolation.

**Graphical Abstract:** 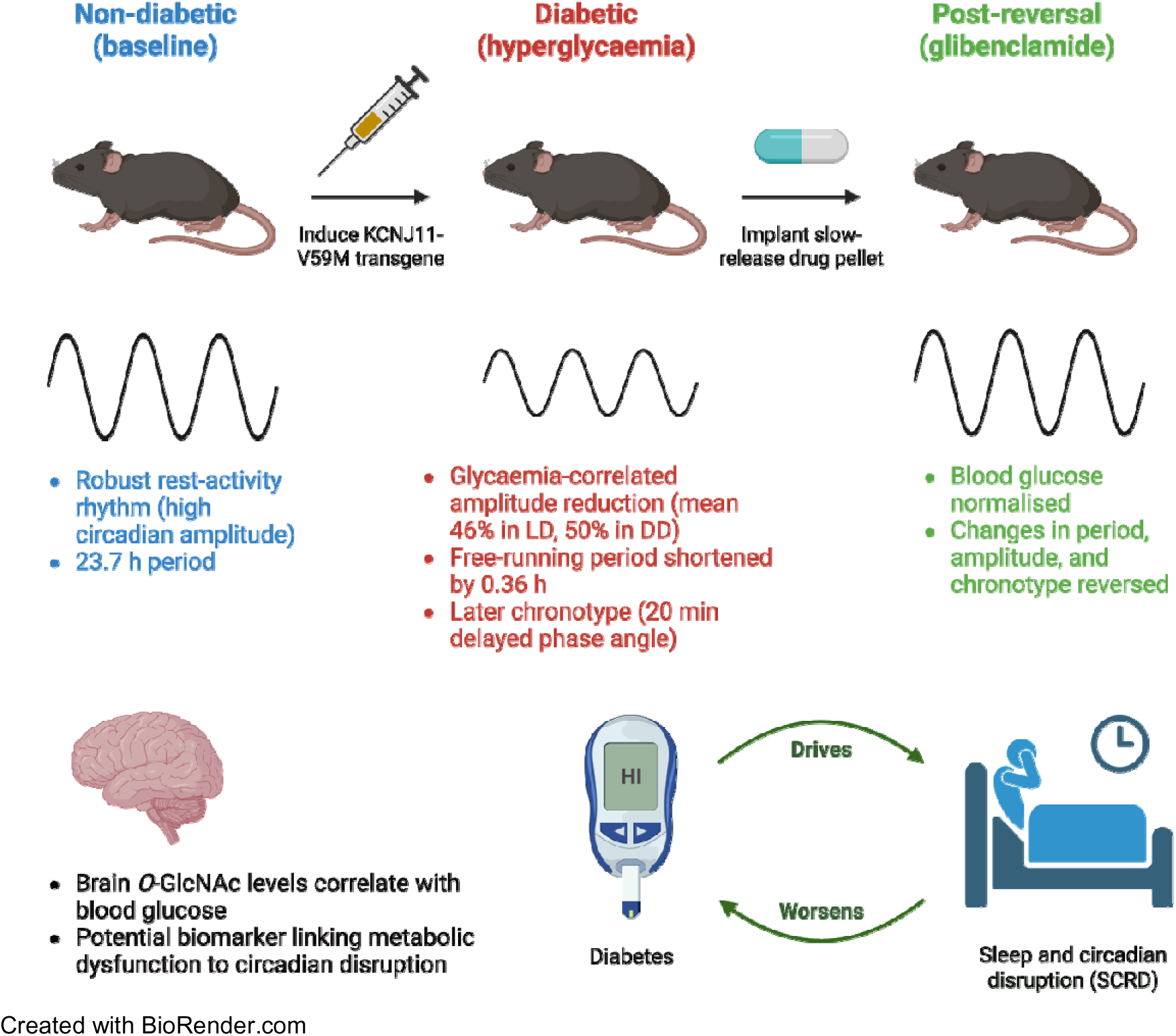

## 1. Introduction

Sleep and circadian rhythm disruption (SCRD) acutely impairs glycaemic control and in the long term increases the risk of developing a range of metabolic disorders including type 2 diabetes (T2D) (Lloyd et al., 2025; Mason et al., 2020; Scheer et al., 2009; Yeung et al., 2025). However, a key unresolved question is whether the opposite case, that diabetes causes SCRD, is also true. Human cohort studies provide correlational evidence that such a relationship may exist but cannot establish directionality, nor separate the effects of hyperglycaemia itself from the comorbidities that accompany it, such as obesity and cardiovascular disease. Impaired glucose tolerance (IGT), indicative of a pre-diabetic state, is associated with dampened circadian amplitude and increased intradaily variability (a measure of rhythm fragmentation) (Xu et al., 2022). Likewise, quantity and quality of sleep predict the risk of developing T2D (Cappuccio et al., 2010), and people with T2D exhibit significantly altered sleep architecture, with less time spent in slow-wave sleep (SWS) and more time in rapid eye movement (REM) sleep (Pallayova et al., 2010). Diabetes is also strongly associated with cognitive impairment and dementia, and bidirectionally associated with mood disorders, with odds of depression being ∼2-fold higher than the general population for the major forms of diabetes (Beverly & Gonzalez, 2025; Biessels & Despa, 2018; Pouwer et al., 2026). SCRD is a common feature connecting these disorders (Cribbet et al., 2016).

We hypothesised that hyperglycaemia-induced circadian disruption could provide a mechanistic link between diabetes and some of these comorbidities. Nutrients are potent circadian zeitgebers (external cues that entrain circadian clocks), especially to peripheral clocks: metabolic cues, such as sudden changes in feeding time, can abruptly shift the phase of peripheral oscillators, uncoupling them from the central pacemaker (Damiola et al., 2000). A high-fat diet extensively remodels the liver clock, with many novel oscillating transcripts and metabolites arising and others lost or phase advanced (Eckel-Mahan et al., 2013). Whether such changes reflect obesity, dyslipidaemia or the accompanying hyperglycaemia has not been resolved, since diet-induced models alter all three together. Meanwhile, the suprachiasmatic nucleus (SCN) – the hypothalamic master clock, comprising ∼10,000 rhythmically firing neurons, entrained by light intensity signals from photosensitive retinal ganglion cells, which co-ordinate local clocks throughout the body via neural, hormonal, and behavioural outputs (Hughes et al., 2015) – contributes to daily rhythms in plasma glucose and glucose uptake and is itself glucose-sensitive, with extracellular glucose altering SCN neuronal firing through ATP-sensitive potassium channels (Hall et al., 1997; Iwanaga et al., 2005; la Fleur, 2003; Yang et al., 2017). The SCN thus integrates metabolic as well as photic signals to co-ordinate brain and peripheral energy status. This raises the possibility that elevated brain glucose could impair central timekeeping, with profound effects on downstream clock-controlled pathways leading to feedback effects that further impair glucose homeostasis. No previous study has longitudinally induced and reversed hyperglycaemia while characterising circadian behaviour within the same animals. Using a model that permits the precise temporal control and reversibility required to do so, we aimed to determine whether diabetes onset can cause SCRD and if this is reversible.

We selected the βV59M mouse model of human neonatal diabetes as this enables rapid and consistent induction of reversible hyperglycaemia and lacks confounding metabolic derangements such as dyslipidaemia (Brereton et al., 2016; Rohm et al., 2018). This model has beta-cell-specific hemizygous expression of the inducible Kir6.2-V59M transgene: when induced, the mutant channels remain open even at high ATP/ADP ratios, preventing beta-cell depolarisation, calcium influx and insulin secretion, thereby causing hypoinsulinaemia and hyperglycaemia (Supplementary Figure 1) (Brereton et al., 2014; Girard et al., 2009). Sulfonylurea drugs such as glibenclamide close the mutant KATP channels and thus reverse diabetes and the associated metabolic dysfunction (Brereton et al., 2014, 2016). We induced and subsequently reversed hyperglycaemia in βV59M mice and tested how rest-activity behaviour was affected, using the pre-induction baseline for each animal as its own control. We examined activity rhythms under a normal 12/12 light/dark cycle (LD) and in constant darkness (DD), the latter allowing endogenous rhythms to be assessed without light entrainment, and tested phase shifting using a simulated jet lag and a light pulse.

O-GlcNAcylation is a nutrient-sensitive protein modification generated through the hexosamine biosynthetic pathway, which integrates glucose availability with amino acid, lipid and nucleotide metabolism (Bouché et al., 2004; Hart et al., 2011). O-GlcNAcylation interacts extensively with phosphorylation (Hart et al., 2011; Kaasik et al., 2013), and core clock proteins including BMAL1, PER2 and CLOCK are rhythmically O-GlcNAcylated, modulating circadian period at cellular and organismal levels (Besing et al., 2015; Li et al., 2013). We therefore asked whether brain O-GlcNAc levels track the hyperglycaemic state *in vivo*, providing a molecular readout of the metabolic environment to which the central clock is exposed.

## 2. Materials and Methods

### 2.1. Animal experiments

All procedures were conducted in accordance with the UK Home Office Animals (Scientific Procedures) Act 1986 and the University of Oxford’s Policy on the Use of Animals in Scientific Research (PPL P3467331), as approved by the local Animal Care and Ethical Review committee (ACER). βV59M mice were generated using a Cre-lox approach on a C57BL6/J background as described previously (Brereton et al., 2014; Girard et al., 2009). Mice of both sexes were used and there was an age difference of 23 days between the oldest and youngest mice in the experimental cohort. The mice were aged 4 months to 9 months over the course of the study. Mice were individually housed in open-top cages (465 cm^2^ floor area) within a light-tight chamber (LTC) equipped with a passive infrared (PIR) sensor above each cage, a validated approach for phenotyping circadian activity and sleep status in mice (Brown et al., 2017). A custom Perspex block was placed under each food hopper to ensure mice remained in range of the PIR sensor at all times. Mice were acclimatised for at least 7 days on a 12/12 light/dark cycle (12 hours of light and 12 hours of dark, henceforth referred to simply as LD) before being subject to altered light/dark cycles. Illuminance was set to 100 lux unless otherwise stated.

At each stage of the study, animals were recorded under LD then subjected to a single 6 h advance of the light/dark cycle, and were recorded under DD, during which they received a 30 min light pulse (50 lux) at approximately CT14, chosen so that all animals received the pulse within the maximal phase delay zone of CT14 to CT16 (Pendergast et al., 2010). Onset of activity on each day was used to measure phase relative to the light/dark cycle. Three animals were culled prior to study completion, two due to surgical complications during pellet implantation, and one due to excessive hyperglycaemia. Exclusions were made on welfare criteria alone, independent of behavioural data.

Transgene expression was induced by subcutaneous injection of 50 mg/kg tamoxifen (Sigma, T5648) dissolved in corn oil (5µl/g body weight) when mice were approximately 6 months old. While all mice in the main study cohort were hemizygous for the Kir6.2-V59M transgene, it has been shown previously that tamoxifen does not affect blood glucose in littermate controls with different genotypes (Brereton et al., 2014). To assess the effect of tamoxifen on rest-activity rhythms, a cohort of wild-type C57BL/6J mice, age-matched to the βV59M cohort, were housed and monitored identically and received tamoxifen at 50 mg/kg. Mice were diabetic for 8 weeks following induction, during which time ALPHA-dri bedding was provided in case of polyuria. After this, a 28-day-release 2mg glibenclamide drug pellet (Innovative Research of America) was implanted subcutaneously using a 10-gauge precision trochar (Innovative Research of America) under isoflurane anaesthesia and the wound was closed using Gluture Topical Tissue Adhesive. Tamoxifen injection and pellet implantation took place within the same five-day windows for all animals. Recordings were made from at least 5 days after induction or pellet implantation (except for ‘diabetes onset’ data, which uses the 5 days starting the day after tamoxifen injection).

Mice had unrestricted access to water and a regular chow diet. Dose and route were as previously established for reversal of hyperglycaemia in this model (Brereton et al., 2014). Body weight was monitored regularly; mice were weighed under red light if LTCs were in darkness. Blood glucose levels were measured from the tail vein at ZT 10-12 during LD using the Freestyle Lite device and Freestyle Lite test strips (both Abbott).

Animals were culled while normoglycaemic (having had their diabetes reversed by glibenclamide therapy) or hyperglycaemic (after the exhaustion of the glibenclamide pellet led to blood glucose increasing again). Animals were culled by cervical dislocation, their terminal blood glucose measurement taken, and tissue samples snap frozen immediately.

### 2.2. Western blotting

Protein was extracted from mouse brain tissue by mechanical homogenisation in RIPA buffer (Sigma-Aldrich, R0278) supplemented with Halt Protease and Phosphatase Inhibitor Cocktail with EDTA added at 1x (Thermo Scientific, 78440). Homogenates were incubated on ice for 30 minutes then centrifuged at 13,000 x g for 15 minutes at 4°C. The protein concentration of the supernatant was determined using a Pierce BCA protein assay kit (Thermo Scientific, 23227). For SDS-PAGE, samples were boiled for 5 minutes in sample loading buffer (LICOR, 928-40004) containing 10% β-mercaptoethanol, and 30µg protein per lane was loaded in a 4–20% Criterion TGX Precast Midi Protein Gel (Bio-Rad, 5671095). For immunoblotting, proteins were transferred onto PVDF membranes using the trans-blot turbo transfer system (Bio-Rad). Membranes were stained with Revert 520 Total Protein Stain (LICOR, 926-10021) for 5 minutes. Membranes were then blocked in Intercept TBS blocking buffer (LICOR, 927-60001) for 1 hour at room temperature. Primary and secondary antibodies were diluted in a 1:1 mixture of Intercept TBS blocking buffer and 1x TBS containing 0.1% tween-20 (TBST). Membranes were incubated overnight at 4°C with 1:2000 anti-*O*-GlcNAc RL2 antibody (Abcam, ab2739). Membranes were washed 3x10 minutes in TBST then incubated with 1:20,000 goat anti-mouse antibody (LICOR, 926-32210) for 1 hour at room temperature. After three additional TBST washes, the membranes were briefly stored in 1x TBS and visualised using the Odyssey Infrared Imaging System (LI-COR Biosciences). Automatic lane finding was used, and signal was normalised to total protein. The immunoblotting protocol was conducted twice and the mean normalised signal for each sample was used for further analysis (membrane scans for each replicate are shown in Supplementary Figure 8).

### 2.3. RT-qPCR

Tissue samples were homogenised in TRIzol reagent using a pestle homogeniser, then shaken vigorously with chloroform and centrifuged at 12,000 x g for 15 minutes at 4°C. The aqueous phase was then used for RNA extraction using the RNeasy mini kit (Qiagen), according to the manufacturer’s instructions. RNA concentration was determined using a NanoDrop spectrophotometer and cDNA was synthesised using a qScript kit (Quantabio, 95047). Quantitative PCR was performed using the Applied Biosystems StepOne Plus Real-Time PCR system. Data were quantified according to the delta-delta Ct method with normalisation to the geometric mean of two housekeeping genes (*Actb* and *Rsp9*). Primer sequences (5’ to 3’) are listed below:

Actb forward: ACCAACTGGGACGATATGGAGAAGA

Actb reverse: CGCACGATTTCCCTCTCAGC

OGT forward: CTGTGTTCGCAGTGACCT

OGT reverse: CAGCCAAATCTCCCCTTGTG

OGA forward: TGCAGTGGTTAGGGTGTCG

OGA reverse: AGCAAACGCTGGAACTCTCC

Rsp9 forward: CTGGACGAGGGCAAGATGAAGC

Rsp9 reverse: TGACGTTGGCGGATGAGCACA

### 2.4. Brain glucose measurements

Following deproteinisation of samples by spin filtering through 3kD MWCO membranes, tissue glucose content was measured using a fluorimetric assay kit (Abcam, ab169559) according to the manufacturer’s instructions. Measured values were normalised to the weight of the tissue sample.

### 2.5. Data processing

PIR raw data in the form of counts per 10 seconds were processed using R and Python. Formatted data were then visualised and analysed using ClockLab software (Actimetrics). In most instances, 7-day windows were used for period and activity analyses. At a minimum, 5-day windows were used where 7 days of suitable data were not available. Analysis windows were selected to avoid gaps in recording, environmental disruptions that impacted activity timing such as cage changes, and periods of acclimatisation and recovery from surgery. Amplitude values were obtained using the ClockLab activity profile function. Period analysis was performed using the Lomb-Scargle and Chi-square methods. Activity onsets were automatically detected by the software but were manually corrected in cases of obvious error. For phase shift analysis, least-squares fit lines were fitted by ClockLab to the daily activity onsets for at least 5 days preceding and following the intervention; phase shift magnitude was calculated as the time difference between the two lines on the day after the intervention. Where data were normalised to overall activity levels, raw counts data for each animal were converted into fractions of total counts in the respective calendar day and then scaled by a factor of 5x10^4^ to convert values to the same order of magnitude as the original data.

### 2.6. Statistical analysis

All statistical analyses were performed in GraphPad Prism [Version 11]. Data are shown as individual values with mean ± SEM, and n refers to the number of animals. Normality was assessed using D’Agostino and Pearson and Shapiro-Wilk tests. Repeated-measures comparisons across study phases were made using a mixed-effects model with the Geisser-Greenhouse correction (to account for missing values, as some animals did not complete all stages of the study), or one-way ANOVA where datasets were complete; non-normally distributed data were analysed by Friedman’s test. Post hoc comparisons used Bonferroni’s or Dunn’s correction as appropriate. Comparisons of two unmatched groups were made using an unpaired two-tailed t-test with Welch’s correction. Relationships between continuous variables were assessed by simple linear regression. The test applied to each comparison is stated in the corresponding figure legend. Differences were considered statistically significant if p < 0.05.

## 3. Results

To determine whether hyperglycaemia disrupts circadian behaviour, and whether any such disruption is reversible, we characterised rest-activity rhythms in the same animals before induction of diabetes, during diabetes, and following reversal of hyperglycaemia (Figure 1). Activity was monitored continuously by PIR throughout, during LD, during entrainment to a simulated jetlag (6h phase advance), during DD, and following a phase-delaying light pulse (30 mins of 50 lux) applied at CT14 (2 hours after activity onset) during DD. To minimise variability, blood glucose was sampled during LD at ZT 10-12 (towards the end of the light phase, when blood glucose levels are least affected by feeding behaviour). Transgene induction increased blood glucose from an average of 6.5 ± 0.25 mM to 23.0 ± 0.56 mM once diabetes was stably established, while glibenclamide treatment subsequently reduced blood glucose to 7.2 ± 0.43 mM post-reversal (Figure 2). Baseline and post-reversal blood glucose levels were slightly higher in males than females, but both sexes reached equal levels of hyperglycaemia (Supplementary Figure 2a).

**Figure 1.**
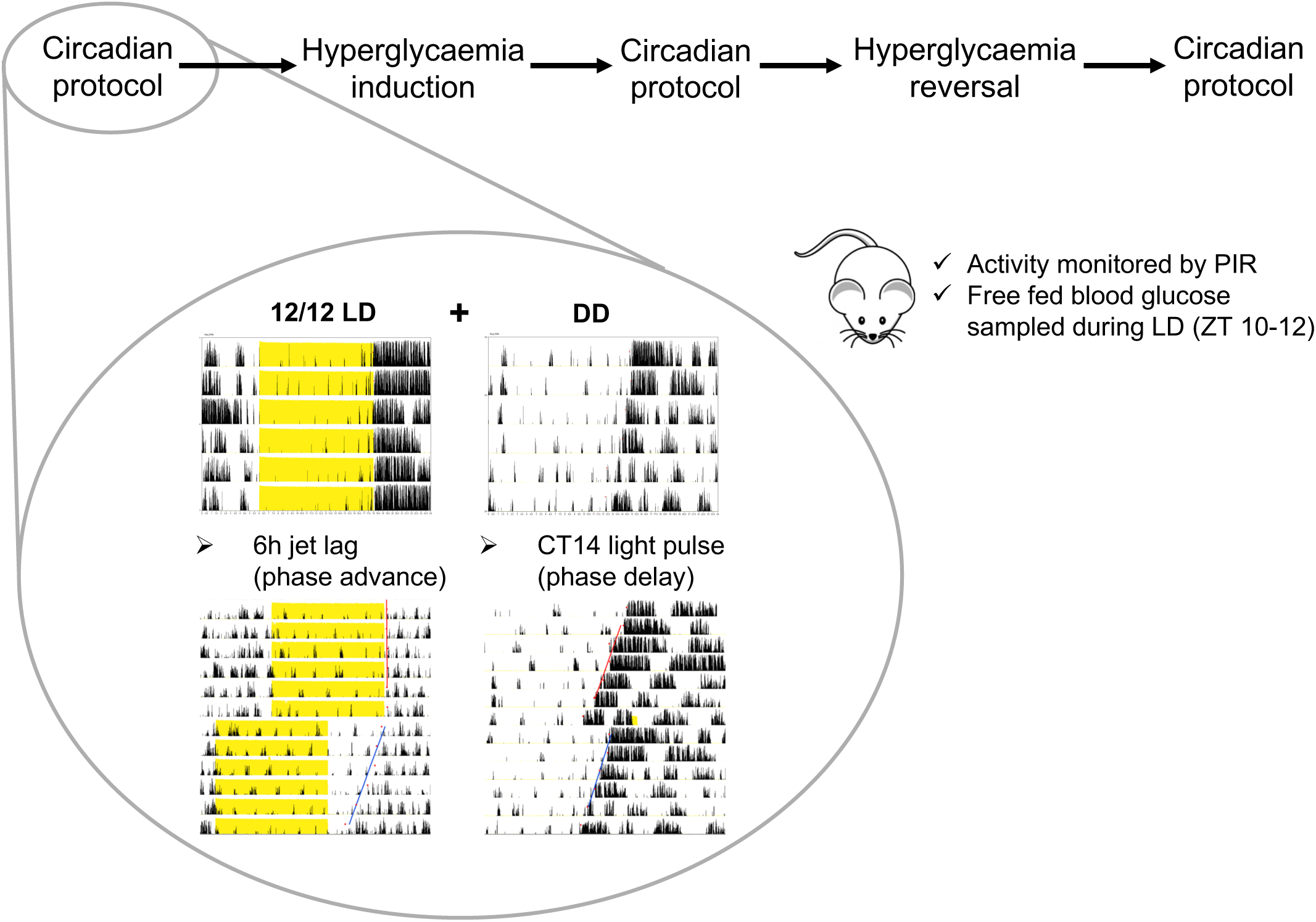
Outline of the design of the study. Animals were subject to a full circadian behavioural characterisation before induction of diabetes (by tamoxifen), while diabetic, and again following reversal of hyperglycaemia (by glibenclamide). This circadian protocol comprised LD, simulated jetlag (6h phase advance), DD, and CT14 light pulse during DD (30 mins of 50 lux). All invasive procedures (including a 1-week recovery period) were performed during LD; during DD, disturbance was kept to a minimum. Throughout the study, mice were subject to continual activity monitoring by PIR, regular weighing, and, during LD, random blood glucose sampling at ZT 10-12. In the representative actograms, which show the activity of an individual animal under different conditions and interventions across (stacked) consecutive days, a yellow background indicates when the lights were on and black spikes indicate activity, with the height of the spike being proportional to intensity (automatically scaled by the ClockLab software). Red dots denote activity onsets, while red and blue lines are fitted to the onsets before and after a simulated jet lag or light pulse.

**Figure 2.**
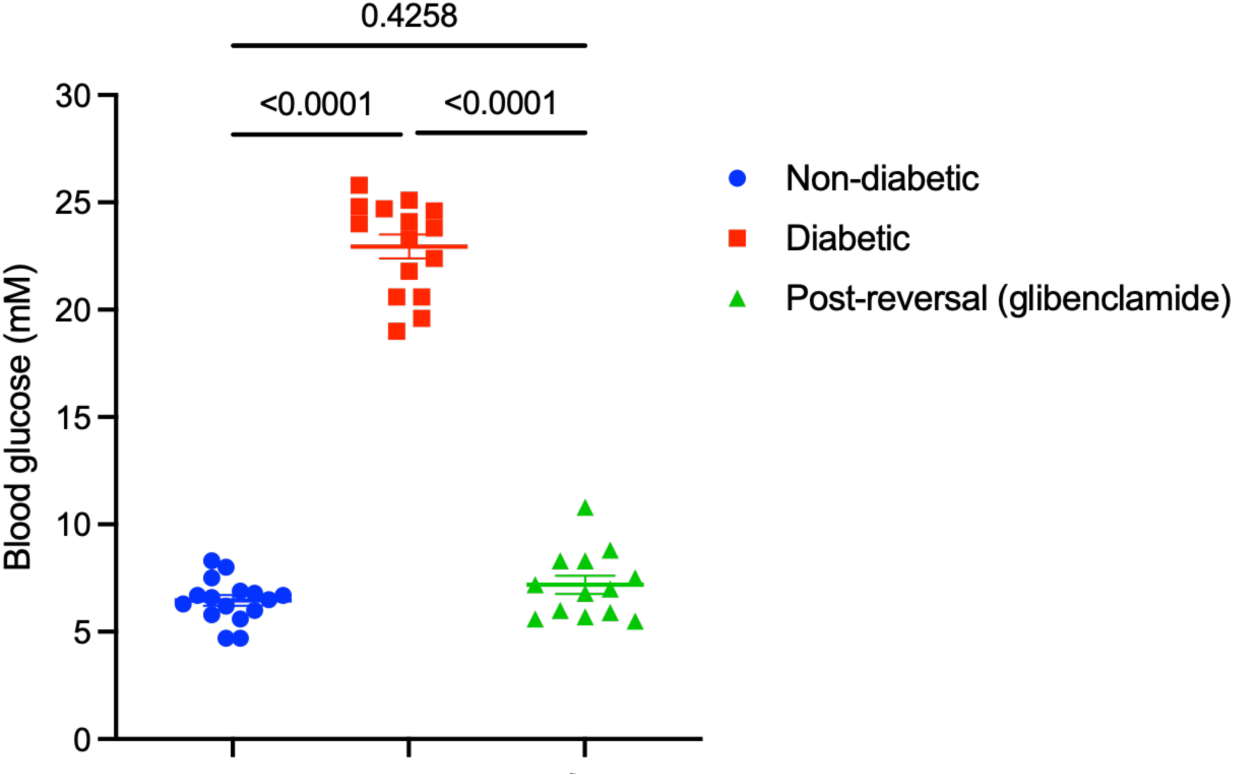
Changes in blood glucose with induction and reversal of hyperglycaemia. Average random blood glucose of mice during each phase of the study: non-diabetic (6.5 ± 0.25 mM, n=16), diabetic (23.0 ± 0.56 mM, n=15), and post-reversal by glibenclamide (7.2 ± 0.43 mM, n=13). Each animal is represented by a single data point per column showing the mean of 2-4 blood glucose readings taken at ZT 10-12 while entrained to a 12:12 LD cycle; error bars display the mean of all animals ± s.e.m. P-values are displayed above (mixed-effects model with Geisser-Greenhouse correction, with Bonferroni’s multiple comparisons test).

### 3.1. Hyperglycaemia reduces the amplitude of rest-activity rhythms

Amplitude in diabetic mice was reduced by an average of 46% in LD and 50% in DD (Figures 3a(i) and 3b(i)). The fall was apparent within days of induction, and the amplitude did not change significantly between the ‘early onset’ and ‘established’ phases of diabetes even though it took almost 2 weeks for blood glucose levels to reach their peak (Figure 3a(iii-iv)). Normalising each animal’s activity to its daily total, thus accounting for changes in overall activity level over time, preserved these effects (Figures 3a(ii) and 3b(ii), and Supplementary Figure 3). The reduction in amplitude is reflected in the daily activity profile, which shows lower peak activity levels and a weaker distinction between the rest and active phases during diabetes (Supplementary Figure 4). In mice with established diabetes, we observed a tight correlation between percentage changes in glycaemia and amplitude (termed Δglucose and Δamplitude): individuals with larger increases in blood glucose showed greater decreases in amplitude (Figures 3a(v) and 3b(iii)). This correlation was strongest for circadian amplitude in constant darkness (r^2^ = 0.60). The amplitude reduction thus appears rapidly once glucose enters the diabetic range, while its magnitude scales with the degree of hyperglycaemia in each animal. In both LD and DD, amplitude did not differ significantly between the non-diabetic and post-reversal groups.

**Figure 3.**
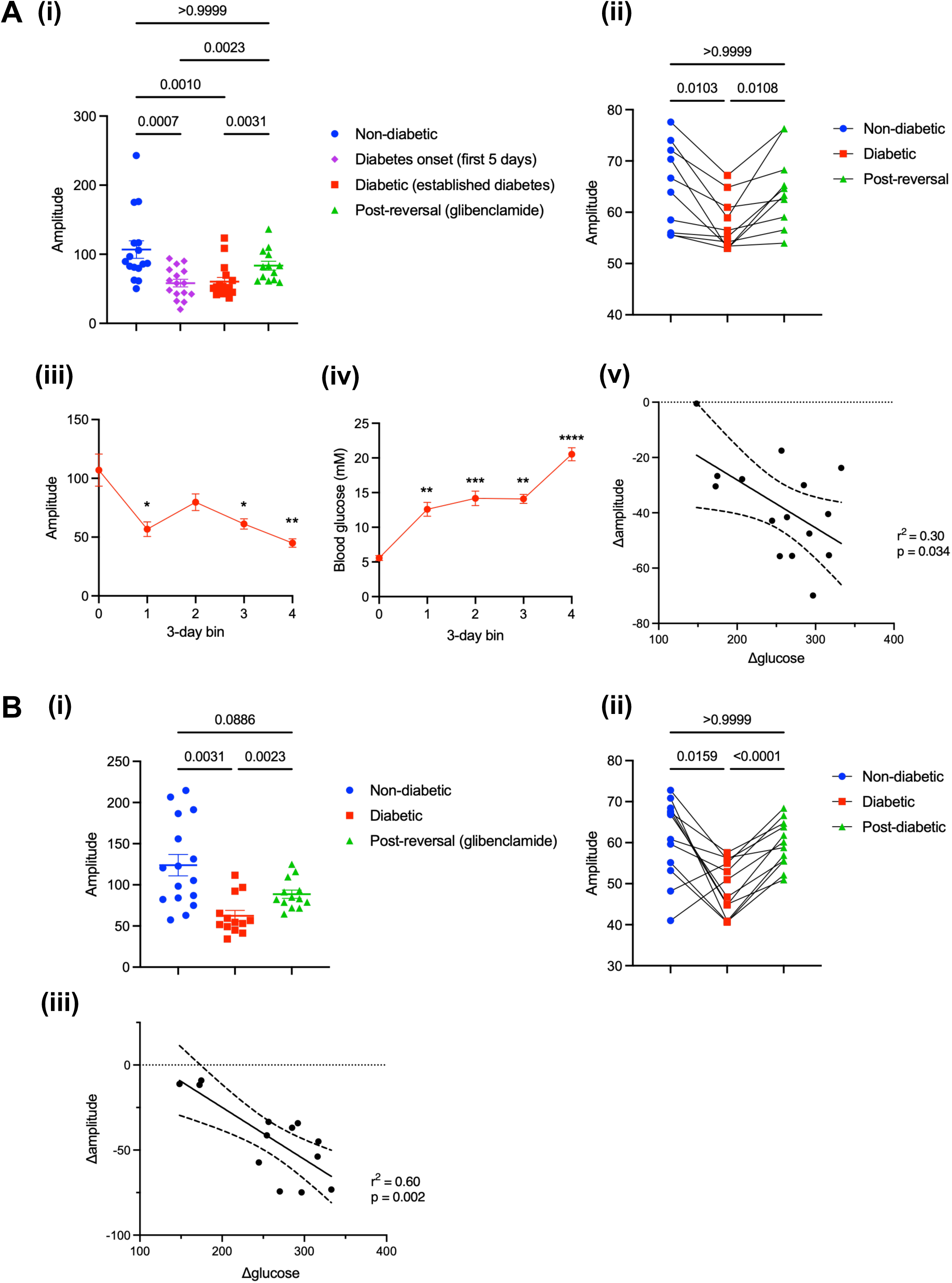
Changes in circadian amplitude following onset and reversal of hyperglycaemia. (a) Amplitude during LD (i) Amplitude analysed during LD at four phases: non-diabetic, diabetes onset (first 5 days following induction), diabetic (established diabetes), and post-reversal (glibenclamide). Individual values are shown for each animal: n=16 (first three phases) or n=13 (post-reversal). Error bars display the mean of all animals ± s.e.m. P-values are displayed above (mixed-effects model with Geisser-Greenhouse correction, with Bonferroni’s multiple comparisons test). (ii) PIR activity data normalised for total daily activity levels: amplitude analysed during LD at baseline, during established diabetes, and following diabetes reversal. Amplitude values displayed with connecting lines to visualise the trajectory of each animal; only complete datasets are included i.e. individuals for which data are available across all phases of the study. Individual values are shown for each animal: n=10. P-values are displayed above (row-matched one-way ANOVA with Bonferroni’s multiple comparisons test). (iii) Circadian amplitude analysed over 3-day windows during the onset of hyperglycaemia, where bin 0 = 3 days before induction, bin 1 = 3 days following induction, and so on. Mean of all animals ± s.e.m. is shown. *p<0.05, **p<0.01 (mixed-effects model with Geisser-Greenhouse correction, with Bonferroni’s multiple comparisons test). (iv) Blood glucose levels during the onset of hyperglycaemia, averaged in 3-day bins as in (iii). Mean of all animals ± s.e.m. is shown. **p<0.01, ***p<0.001, ****p<0.0001 (Friedman test with Dunn’s multiple comparisons test). (v) Percentage change in circadian amplitude following onset of hyperglycaemia (Δamplitude) plotted against % change in blood glucose following onset of hyperglycaemia (Δglucose). Simple linear regression with 95% confidence intervals indicated by dashed lines; r^2^ = 0.30, p = 0.034 for deviation of slope from zero. (b) Circadian amplitude during DD (i) Circadian amplitude during DD at three phases: non-diabetic, diabetic (established diabetes), and post-reversal (glibenclamide). Individual values are shown for each animal: n=16 (non-diabetic) or n=13 (diabetic and post-reversal). Error bars display the mean of all animals ± s.e.m. P-values are displayed above (mixed-effects model with Geisser-Greenhouse correction, with Bonferroni’s multiple comparisons test). (ii) PIR activity data normalised for total daily activity levels: amplitude analysed during LD at baseline, during established diabetes, and following diabetes reversal. Amplitude values displayed with connecting lines to visualise the trajectory of each animal; only complete datasets are included i.e. individuals for which data are available across all phases of the study. Individual values are shown for each animal: n=12. P-values are displayed above (row-matched one-way ANOVA with Bonferroni’s multiple comparisons test). (iii) Percentage change in circadian amplitude following onset of hyperglycaemia (Δamplitude) plotted against % change in blood glucose following onset of hyperglycaemia (Δglucose). Simple linear regression with 95% confidence intervals indicated by dashed lines; r^2^ = 0.60, p = 0.002 for deviation of slope from zero.

To test whether tamoxifen itself affected rest-activity rhythms independently of transgene induction, we monitored wild-type mice by PIR before, during, and after tamoxifen dosing (50 mg/kg, during LD, age-matched to the βV59M mice). Amplitude fell transiently during dosing (p = 0.0476) and returned fully to baseline within 3 days (Supplementary Figure 5). Amplitude in βV59M mice remained reduced for the full 8 weeks of diabetes and recovered only on restoration of euglycaemia.

To assess the consistency of an individual’s rest-activity rhythms, we analysed intradaily variability (IV, fragmentation within days) and interdaily stability (IS, day-to-day regularity) during LD and DD (Van Someren et al., 1999; Witting et al., 1990; Xu et al., 2022). Neither changed significantly during diabetes, other than a transient rise in IS during diabetes onset (Supplementary Figure 6). The reduction in amplitude therefore reflects a loss of day-night contrast rather than fragmentation or reduced interdaily stability.

### 3.2. Hyperglycaemia shortens free-running period in constant darkness

To determine whether hyperglycaemia affects the endogenous clock rather than its entrainment to light, we measured free-running period in constant darkness. Period was unchanged under 12/12 LD, as expected in entrained animals (Figure 4a), but in DD diabetic mice showed a shortening of 0.36 h that reversed on restoration of euglycaemia (Figure 4b). Both Lomb-Scargle (p = 0.0036, Figure 4b(i)) and Chi-square (p = 0.0019, data not shown) analyses showed significant shortening. Period fell from 23.69 h to 23.33 h, the baseline being consistent with published values for C57BL/6J mice (Gutman et al., 2011), and the resulting faster daily advance of activity onset is visible in representative actograms from the same individual at each phase of the study (Supplementary Figure 7a).

**Figure 4.**
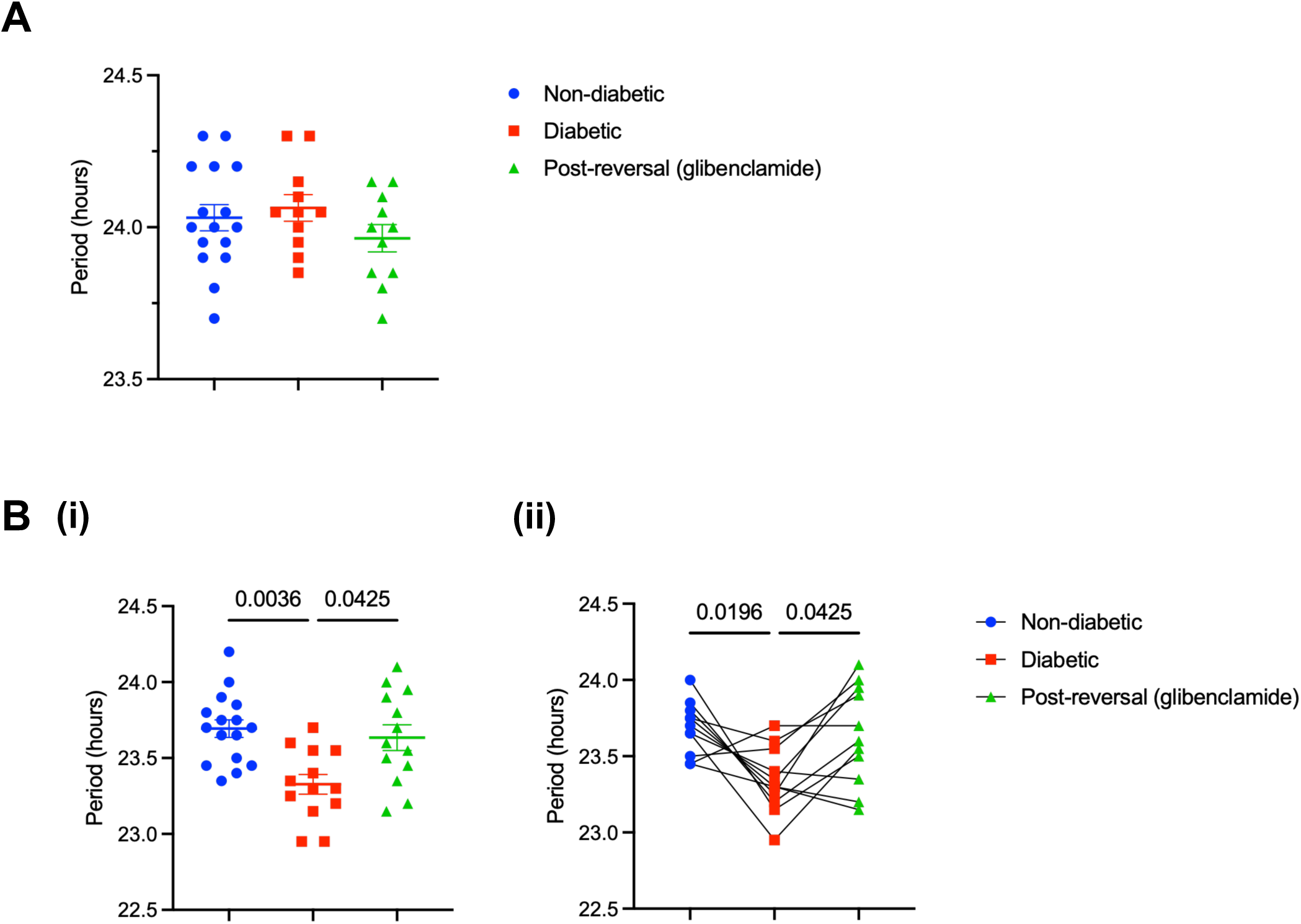
Changes in period following onset and reversal of hyperglycaemia. Circadian period calculated using the Lomb-Scargle method during (a) LD and (b) DD at three phases: non-diabetic, diabetic, and post-reversal (glibenclamide). Part b(ii) shows period values with connecting lines to visualise the trajectory of each animal; only complete datasets are included i.e. individuals for which data are available across all phases of the study. Individual values are shown for each animal: n=16 (non-diabetic) or n=13 (diabetic and post-reversal), except for part b(ii), where n=8. Error bars display the mean of all animals ± s.e.m. P-values are displayed above (mixed-effects model with Geisser-Greenhouse correction or row-matched one-way ANOVA (b(ii)) with Bonferroni’s multiple comparisons test).

### 3.3 Hyperglycaemia delays chronotype and slows re-entrainment without altering the phase-delay response to light

We calculated the phase angle of entrainment (the interval between activity onset and lights off) for each mouse during LD. We observed a difference of 20 minutes in phase angle: whereas non-diabetic mice anticipated lights off, diabetic mice did not become active until after the lights turned off (Figure 5a), indicating a delayed chronotype.

**Figure 5.**
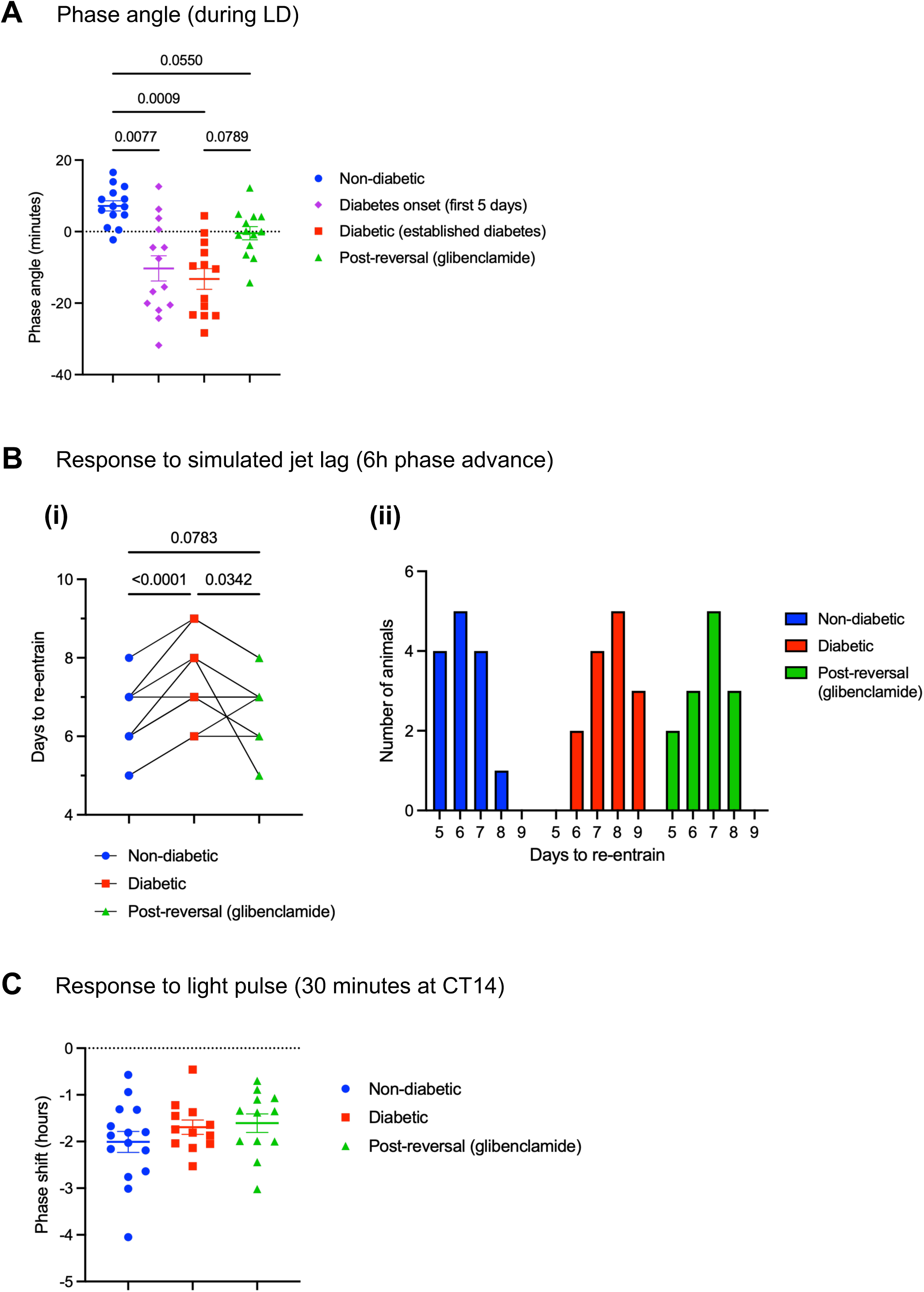
Changes in phase shifting following onset and reversal of hyperglycaemia. (a) Phase angle (in minutes) during LD at four phases: non-diabetic, diabetes onset (first 5 days following induction), diabetic (established diabetes), and post-reversal (glibenclamide). Phase angle is calculated as the time difference between activity onset and lights off: the value is positive when the onset precedes lights off and negative when the onset occurs after lights off. P-values are displayed above (mixed-effects model with Geisser-Greenhouse correction, with Bonferroni’s multiple comparisons test). (b) Re-entrainment following a 6-hour jet lag (lights on after 6h of darkness) at three study phases: non-diabetic, diabetic, and post-reversal (glibenclamide). Days to re-entrain is the number of days for activity onsets to re-align with lights off (based on slope fitted to onsets starting at day 1 following the jet lag). (i) Individual values are shown for each animal with connecting lines to visualise individual trajectories through the three study phases: n=14 (non-diabetic and diabetic) or n=13 (post-reversal). P-values are displayed above (mixed-effects model with Geisser-Greenhouse correction, with Bonferroni’s multiple comparisons test). (ii) Same data as (i), presented as a frequency distribution. (c) Phase shift (in hours) following a light pulse (30 mins of 50 lux) at approximately CT14 at three phases: non-diabetic, diabetic, and post-reversal (glibenclamide). Individual values are shown for each animal; error bars display the mean of all animals ± s.e.m. P-values are displayed above (mixed-effects model with Geisser-Greenhouse correction, with Bonferroni’s multiple comparisons test).

To determine whether hyperglycaemia alters the way the clock responds to light, we investigated phase shifting using two paradigms, one phase-advancing (6 h simulated advancing jet lag) and one phase-delaying (light pulse).

Following a 6 h phase advance, diabetic mice took, on average, 36 hours longer than non-diabetic mice, and 23 hours longer than post-reversal mice, to re-entrain to the new light/dark cycle (Figure 5b). Representative actograms are shown in Supplementary Figure 7b.

For the light pulse intervention, we aimed to deliver light at CT14, i.e. 2 hours after activity onset, within the maximal phase delay zone of CT14 to CT16 (Pendergast et al., 2010). Because individual phases of mice diverged in DD, pulses were delivered at CT13.7 ± 0.9 h SD for non-diabetic mice, CT14.9 ± 0.9 h SD for diabetic mice, and CT14.3 ± 1.0 h SD for post-reversal mice. The magnitude of phase delay did not differ between study phases (Figure 5c), indicating that the phase-delay response is unaltered by hyperglycaemia. Representative actograms are shown in Supplementary Figure 7c.

### 3.4 Brain O-GlcNAcylation tracks blood glucose in diabetes

To investigate whether brain protein O-GlcNAcylation tracks the prevailing glucose concentration and could therefore report on the altered glycaemic state of the diabetic brain, we quantified O-GlcNAc in the whole brain by immunoblotting. We found a significant positive correlation between whole-brain O-GlcNAc levels and terminal blood glucose (r² = 0.57, p = 0.002; Figure 6a; original images shown in Supplementary Figure 8). Individuals were stratified by terminal blood glucose, as either normoglycaemic or hyperglycaemic according to an established threshold of 12 mM (Haythorne et al., 2026), reflecting whether glycaemia was still corrected by glibenclamide at the time of cull (see Methods, section 2.1). O-GlcNAc was significantly elevated in the hyperglycaemic group (Figure 6b). Paralleling this, whole-brain glucose content was also significantly elevated in hyperglycaemia (Supplementary Figure 9), consistent with increased glucose availability in the brain.

**Figure 6.**
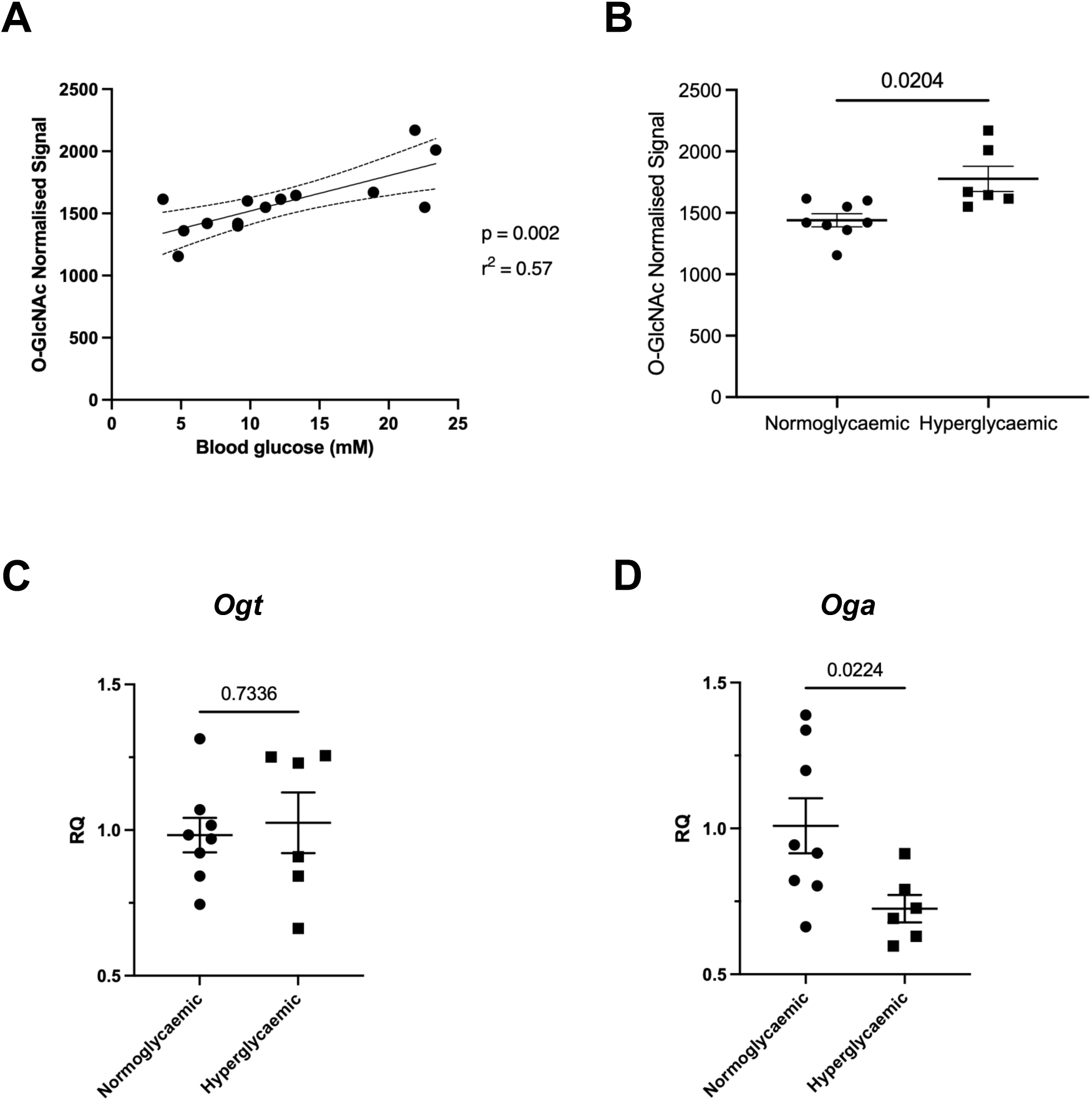
Analysis of brain O-GlcNAc levels and expression of OGT and OGA. (a) Brain O-GlcNAc levels plotted against terminal blood glucose. Western blot signal is normalised to total protein stain. Simple linear regression with 95% confidence intervals indicated by dashed lines; r^2^ = 0.57, p = 0.002 for deviation of slope from zero. (b) Same data as in (a), grouped according to blood glucose: normoglycaemic (≤12 mM, n = 8) or hyperglycaemic (>12 mM, n = 6). P-value displayed above (unpaired two-tailed t-test with Welch’s correction). (c,d) Relative mRNA levels of the genes (c) *Ogt* and (d) *Oga* in brain lysates from normoglycaemic (terminal blood glucose ≤12 mM, n=8) and hyperglycaemic mice (terminal blood glucose >12 mM, n=6). P-value displayed above (unpaired two-tailed t-test with Welch’s correction).

The addition and removal of O-GlcNAc is catalysed by O-GlcNAc transferase (OGT) and O-GlcNAcase (OGA) respectively (Hart et al., 2011). To investigate the mechanism underlying increased O-linked glycosylation in hyperglycaemia, we measured the expression of these genes. Relative to normoglycaemic brains, *Ogt* was unchanged in hyperglycaemia (Figure 6c), whereas *Oga* was significantly downregulated in hyperglycaemic brains (Figure 6d). If these changes in gene expression are reflected at the level of enzyme activity, downregulation of OGA without any change in OGT would be expected to cause O-GlcNAc levels to increase over time due to the slower off rate for the modification.

## 4. Discussion

### 4.1 Changes in amplitude and period

We observed a striking drop in the amplitude of rest-activity rhythms in diabetic mice, the magnitude of which corresponded to the increase in blood glucose (Figure 3). These findings remained robust even after accounting for variation in daily activity (Figures 3a(ii) and 3b(ii), and Supplementary Figure 3). The decline in amplitude was deeper and more sustained than could be explained by any acute effect of tamoxifen (Supplementary Figure 5), persisting throughout the 8 weeks of diabetes, and was reversed only after restoration of euglycaemia. A comparable reduction in nocturnal activity amplitude has been reported in 16-week high-fat-diet C57BL/6J mice (Cunningham et al., 2016), indicating that reduced rest-activity amplitude is a consistent feature of diabetic states across models, although the multiple metabolic changes in diet-induced models make the contribution of glycaemia difficult to isolate compared to our model where hyperglycaemia is induced without obesity, dyslipidaemia or insulin resistance.

Diabetic mice also showed a shortening of free-running period, an effect masked during LD by entrainment and revealed only in DD. Sinturel et al. (2019) first linked glycaemic state to clock speed in humans, showing an inverse correlation between HbA1c and cellular circadian period in fibroblasts from people with T2D. By inducing and then reversing hyperglycaemia within the same animals, we now show that this relationship is causal and reversible *in vivo.* Direct effects of glucose on central clock neurons offer a possible route: increased glucose availability shortens the period and delays the phase of Per2 expression in hypothalamic neurons (Oosterman & Belsham, 2016), mirroring both changes observed here and consistent with the elevated brain glucose in our diabetic animals (Supplementary Figure 9). In line with this, SCN Per2 expression changes in response to glucose *in vivo* (Iwanaga et al., 2005), and SCN neurons are themselves intrinsically glucose-sensitive, with extracellular glucose altering their circadian firing pattern (Hall et al., 1997).

### 4.2 Changes in chronotype and re-entrainment kinetics

Our phase-shifting results are intriguing because they do not follow from the changes in period and amplitude. A shorter endogenous period leaves an oscillator already drifting in the direction of an advancing shift, and a lower-amplitude oscillator is more readily reset by a given stimulus (Abraham et al., 2010); both therefore predict faster re-entrainment to a phase advance and an earlier phase angle of entrainment. Diabetic animals showed the opposite on both counts. Thus, phase-advancing behaviour was impaired despite the changes in period and amplitude, while the phase-delay response to light was unaltered.

Together, these findings point to an additional effect of hyperglycaemia on the phase-shifting properties of the circadian system, the nature of which will require further investigation. We measured delays directly but infer the advance impairment from re-entrainment; a full phase response curve would be needed to distinguish reduced advance responsiveness from altered phase dependence. Glycaemic state has previously been shown to modulate photic phase shifting, with light-induced phase delays enhanced in streptozotocin-diabetic mice, in which lowering glucose with insulin restored them (Challet et al., 1999). However, as streptozotocin induces diabetes through β-cell destruction and is associated with broader metabolic abnormalities and altered brain signalling (Gelling et al., 2006; Willecke et al., 2015), the resulting diabetic state is not directly comparable to ours.

Evening chronotype is consistently associated with type 2 diabetes and reduced insulin sensitivity in human cohorts, with evening types carrying 25% higher odds of T2D than morning types in the UK Biobank (Tan et al., 2020), and this association holding for meta-analysis (Zhang et al., 2022). The association is usually interpreted as chronotype modifying diabetes risk. Our finding that hyperglycaemia delays chronotype, and that this reverses on restoration of euglycaemia, shows that causation can also run from glycaemia to chronotype.

The effect of glycaemia on amplitude and period appeared more pronounced in females than in males (Supplementary Figure 2), despite no significant sex difference in the change in blood glucose following induction (Supplementary Figure 2a(ii)). The difference resolved on restoration of euglycaemia, indicating a state-dependent effect rather than a constitutional difference between sexes. The study was not powered to test for sex differences, and this observation warrants dedicated investigation.

### 4.3 Brain O-GlcNAc and hyperglycaemia

The correlation of brain O-linked glycosylation with blood glucose indicates that O-GlcNAc tracks the altered glycaemic state of the diabetic brain. This is consistent with the role O-GlcNAcylation plays as a metabolic sensor for clock regulation, rhythmically modifying BMAL1, PER2 and CLOCK co-ordinately with, and in competition with, phosphorylation (Li et al., 2013). The downregulation of *Oga* we observe offers a more likely explanation for the rise in O-GlcNAc than increased hexosamine flux alone. Substrate availability does not by itself determine O-GlcNAc levels: in Alzheimer’s disease models, brain O-GlcNAcylation is reduced despite elevated UDP-GlcNAc (Hawkinson et al., 2026). Whether this sustained rise contributes to the circadian phenotype or merely reflects the metabolic state that produces it remains to be established; either way, it marks a genuine shift in the brain’s metabolic environment that warrants direct interrogation. Resolving circadian variation in brain O-GlcNAc would be informative, particularly during sleep-wake transitions, which coincide with the two daily peaks in synaptic phosphorylation (Brüning et al., 2019).

### 4.4 Strengths and limitations

The βV59M mouse provides a clean, inducible and reversible model of hyperglycaemia, without obesity or dyslipidaemia (Brereton et al., 2016; Rohm et al., 2018). It carries a human *KCNJ11* mutation that causes permanent neonatal diabetes, and like these patients the mice respond to sulfonylurea therapy. The hypoinsulinaemic hyperglycaemia is characteristic of both K_ATP_-dependent neonatal diabetes and T1D. Insulin insufficiency also develops during T2D progression, although in this model, unlike in T2D, insulin sensitivity remains intact (Brereton et al., 2014) and insulin secretion is abrogated more abruptly than in T2D. Expression of the mutant channel is restricted to the beta cell, so the circadian changes we observe cannot be attributed to altered K_ATP_ channel function in neurons. The same model has, however, previously been used to attribute changes in islet structure and function to hyperglycaemia rather than to K_ATP_ channel activation, on the grounds that the changes are reversed by lowering glucose using insulin (Brereton et al., 2014), and here the graded relationship between glycaemia and amplitude, together with full reversal on restoration of euglycaemia, points to glucose as the operative variable. Reversal was achieved pharmacologically, but glibenclamide administered by subcutaneous pellet does not reach detectable concentrations in brain or cerebrospinal fluid of mice despite high plasma levels (Lahmann et al., 2015) so the recovery reflects normalisation of blood glucose rather than a central action of the drug.

Reduced locomotor activity accompanies many illnesses and could, in principle, contribute to these changes. Nevertheless, normalisation to daily total activity preserved the effects, indicating a reduction in the day-night contrast of activity rather than in its overall amount. Moreover, the fall in amplitude was established before glucose peaked; and both amplitude and period recovered on restoration of euglycaemia. A non-specific consequence of ill health would be expected to alter the amount of activity rather than to reinstate the period and amplitude structure of the rhythm.

The within-subjects design has the major benefit of each animal being its own control, controlling for heterogeneity in circadian behaviour between individuals. Animals necessarily age over the course of a longitudinal study, but age increases monotonically across the protocol whereas circadian amplitude and period fall during hyperglycaemia and recover following its reversal, a pattern that ageing cannot produce.

While PIR provides a validated measure of rest-activity rhythms, with immobility scoring only slightly overestimating total sleep time relative to EEG-defined sleep (Brown et al., 2017), we did not directly measure sleep architecture in this study. EEG/EMG telemetry would allow us to interrogate how sleep architecture changes with rapid onset and reversal of hyperglycaemia. Brain O-GlcNAc was measured in whole tissue at a single time point at cull, several weeks after the circadian recordings were made, so we could not test whether O-GlcNAc levels correlate with the circadian phenotype of individual animals.

Because insulin secretion and glycaemia are corrected together in this model, their contributions cannot be separated by reversal alone. Established insulin-to-clock mechanisms, however, do not readily predict the phenotype observed here. Insulin drives PERIOD synthesis and resets peripheral clocks, yet direct insulin treatment of the intact SCN alters neither phase nor period (Crosby et al., 2019). Moreover, insulin-PI3K-AKT signalling inhibits FOXO3, a positive regulator of Clock expression and circadian amplitude, such that reduced insulin signalling would predict higher rather than lower amplitude through this pathway (Chaves et al., 2014). A contribution from altered insulin signalling cannot be excluded, but these observations, together with elevated brain glucose and the direct effects of glucose on hypothalamic clock gene expression and SCN neuronal activity in preparations where insulin is absent (Hall et al., 1997; Oosterman & Belsham, 2016), make glucose the more parsimonious explanation for the central circadian phenotype.

### 4.5 Conclusions

Our results demonstrate that hyperglycaemia can disrupt circadian rest-activity behaviour, in proportion to the severity of hyperglycaemia and reversibly on normalisation of blood glucose. Considering this discovery together with existing evidence, there is clearly a mechanistic basis for considering glucose dysregulation and circadian disruption as reciprocal processes in diabetes, with implications for T2D aetiology and disorders of the brain. If the relationship holds in humans, it argues that both should be managed together: stabilising glycaemia may help protect circadian and sleep-wake function, while stabilising circadian and sleep-wake behaviour may in turn support glycaemic control. Given the high incidence of psychiatric and neurodegenerative disorders in patients with diabetes, and the roughly two-fold higher prevalence of depression (Beverly & Gonzalez, 2025), this relationship demands further exploration as both a mechanism and a treatment target. Even small reductions in circadian amplitude are significantly associated with impaired glucose homeostasis in a general population of adults without diabetes (Xu et al., 2022), and elevations in fasting glucose as small as 3 mM can impair beta-cell function (Haythorne et al., 2026). Sleep delay impairs glycaemic control through changes in SCN neuronal activity, an effect corrected by recovery sleep (Hurtado-Alvarado et al., 2023). Our finding that hyperglycaemia in turn disrupts SCN-driven rest-activity behaviour places the central clock on both arms of this relationship, raising the possibility that small changes in circadian timing and glucose homeostasis exacerbate one another during progression from IGT to T2D.

## Abbreviations

AD: Alzheimer’s disease
BMAL1: Brain and muscle ARNT-like 1
βV59M mouse: mouse model with beta-cell-specific expression of a K_ir_6.2-V59M transgene
CT: circadian time (activity onset occurs at CT12)
DD: dark/dark (constant darkness)
HbA1c: glycated haemoglobin
HBP: hexosamine biosynthetic pathway
HFD: high-fat diet
IGT: impaired glucose tolerance
IS: interdaily stability
IV: intradaily variability
K_ATP_ channel: ATP-sensitive potassium channel
K_ir_6.2/KCNJ11: potassium voltage-gated channel subfamily J member 11
LD: light/dark (12/12 cycle)
LTC: light-tight chamber
*O*-GlcNAc: *O*-linked N-acetylglucosamine
PIR: passive infrared
REM: rapid eye movement
SCN: suprachiasmatic nucleus
SCRD: sleep and circadian rhythm disruption
SWS: slow-wave sleep
T2D: type 2 diabetes
UDP-GlcNAc: uridine diphosphate N-acetylglucosamine
ZT: Zeitgeber time (lights turn on at ZT0)

## Supplementary Material

This article contains supplementary material.

## CRediT authorship contribution statement

Matthew Lloyd: Conceptualisation, Methodology, Investigation, Formal analysis, Writing – Original draft preparation, Writing – Reviewing and Editing, Visualisation.

Guohao Ni: Methodology, Investigation, Writing – Reviewing and Editing

Kai-Ting Hu: Investigation

Russell G Foster: Methodology, Writing – Reviewing and Editing

Frances M Ashcroft: Resources, Writing – Reviewing and Editing

Aarti Jagannath: Methodology, Writing – Reviewing and Editing, Supervision, Project administration, Funding acquisition.

Sridhar R Vasudevan: Conceptualisation, Methodology, Investigation, Writing – Reviewing and Editing, Supervision, Project administration, Funding acquisition for the project.

## Acknowledgments

M.L. wishes to thank Elizabeth Haythorne (University of Edinburgh) for invaluable advice relating to the βV59M mouse model and Manu Verma (Novo Nordisk) for serving as his mentor during his postdoctoral fellowship.

All authors would like to thank Simona Di Pretoro and the animal facility staff, especially Angela Curran and Laura Thomas, for animal care and practical support throughout the study.

