## Supplementary figures for "Hyperglycaemia reversibly disrupts circadian behaviour in mice"

### Supplementary Figure 1

βV59M mouse model:
Tamoxifen-induced expression of mutant Kir6.2 in beta-cells prevents insulin secretion, causing hyperglycaemia, reversible by sulfonylurea therapy

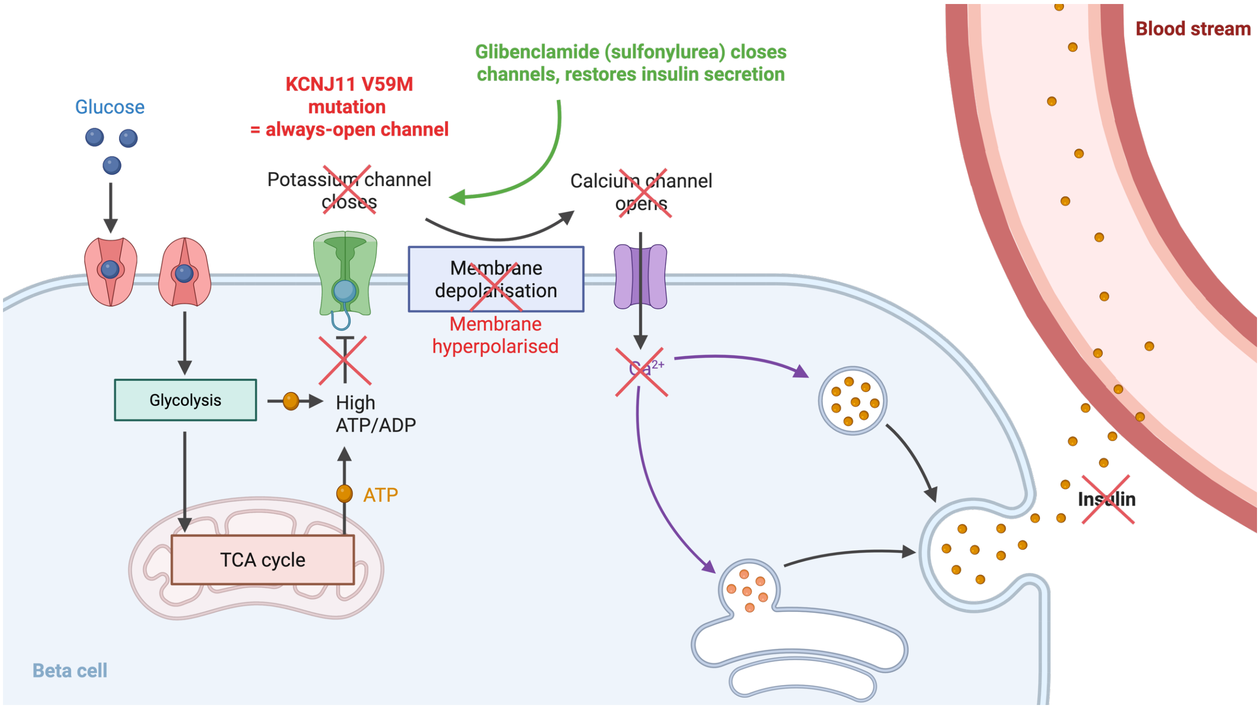


Created with BioRender.com

### Supplementary Figure 2

Data stratified by sex:
(A) Blood glucose, (B) Amplitude in LD, (C) Amplitude in DD, (D) Period in DD

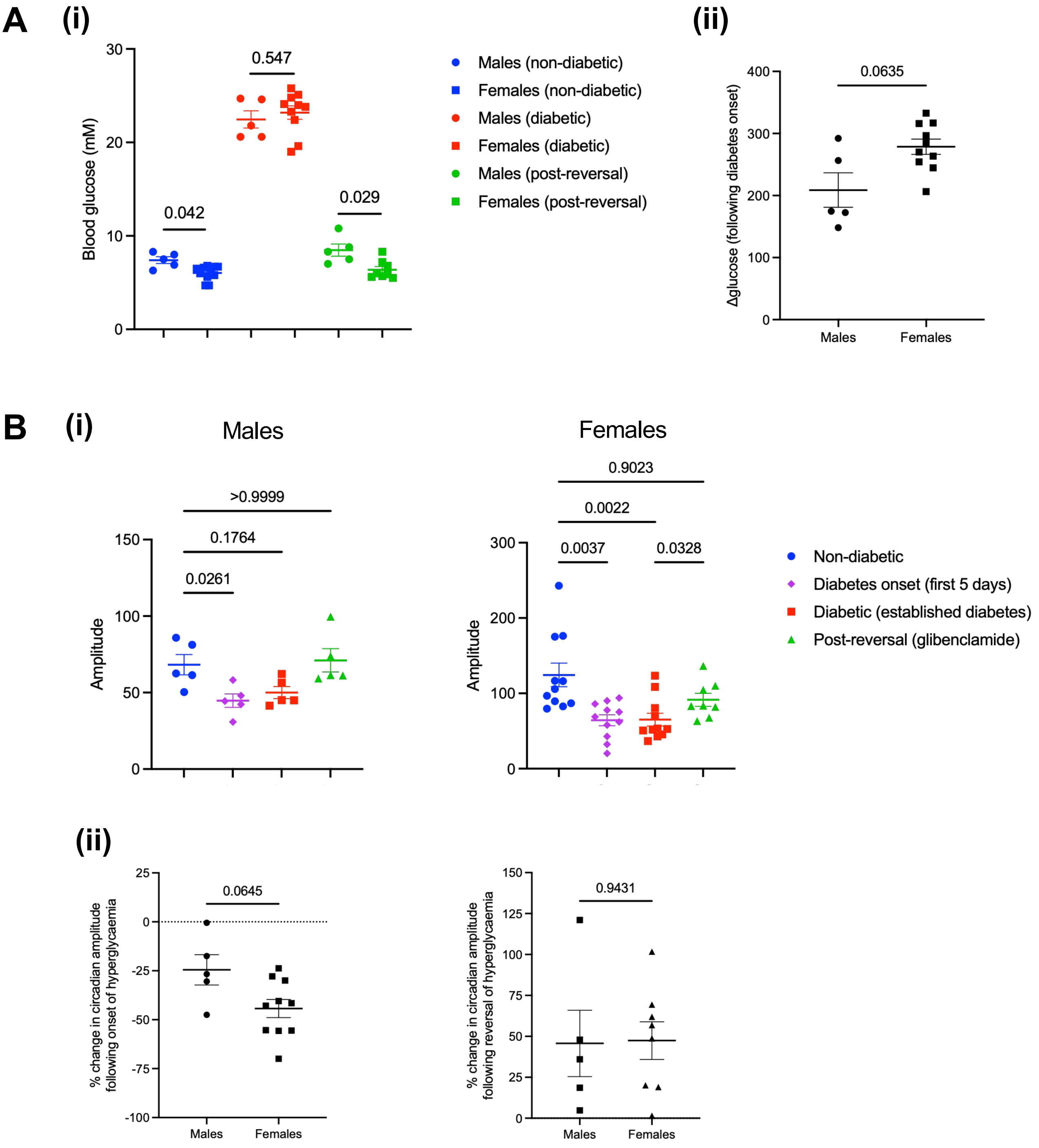

**C**

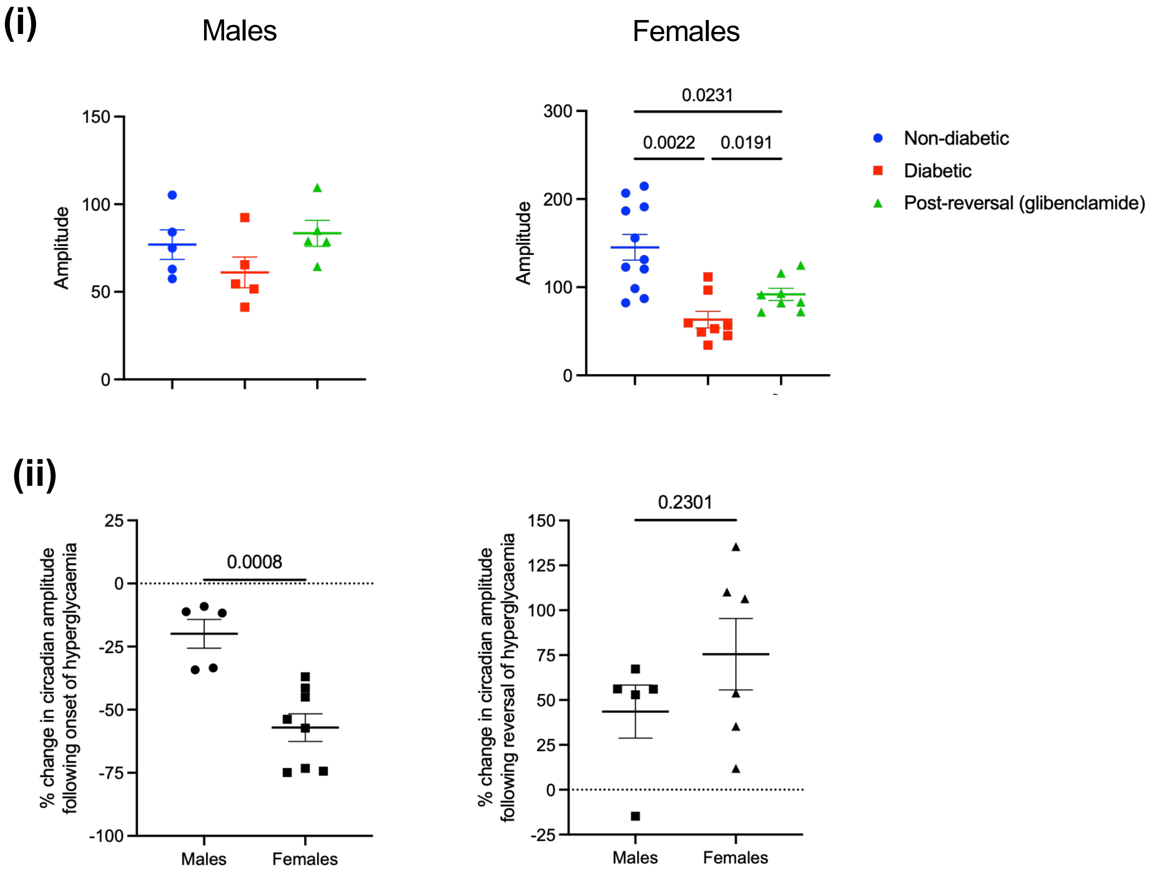
(i)
Males
150
100
50
0
Amplitude
Females
0.0231
0.0022
0.0191
300
200
100
0
Amplitude
Non-diabetic
Diabetic
Post-reversal (glibenclamide)
(ii)
25
0
-25
-50
-75
-100
0.0008
% change in circadian amplitude following onset of hyperglycaemia
Males
Females
0.2301
150
125
100
75
50
25
0
-25
% change in circadian amplitude following reversal of hyperglycaemia
Males
Females

**D**

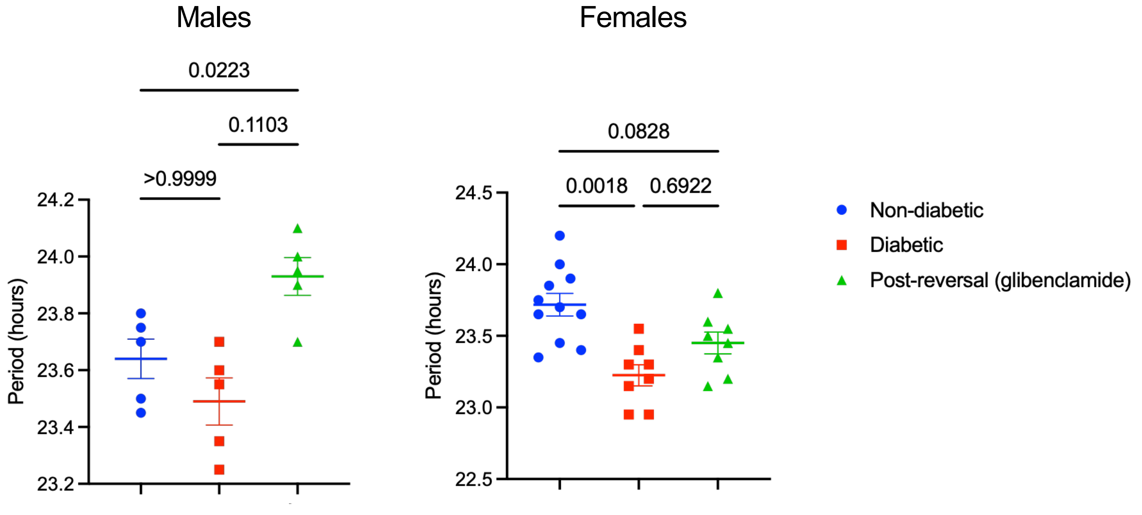
Males
0.0223
0.1103
>0.9999
24.2
24.0
23.8
23.6
23.4
23.2
Period (hours)
Females
0.0828
0.0018
0.6922
24.5
24.0
23.5
23.0
22.5
Period (hours)
Non-diabetic
Diabetic
Post-reversal (glibenclamide)

### Supplementary Figure 3

Changes in circadian amplitude following onset and reversal of hyperglycaemia in (A) LD and (B) DD: PIR activity data normalised for total daily activity levels

**A** (i)

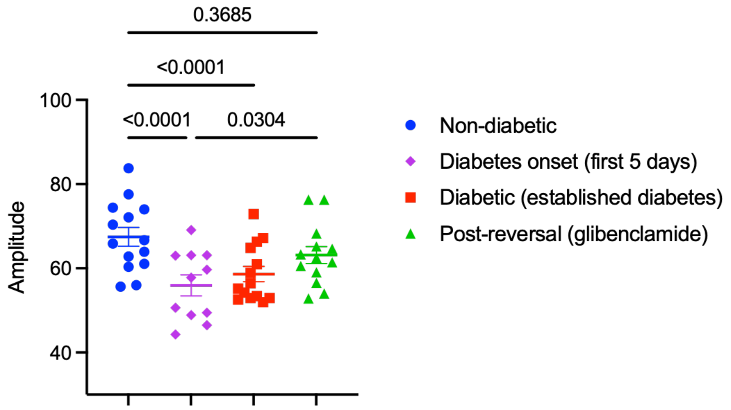


(ii)

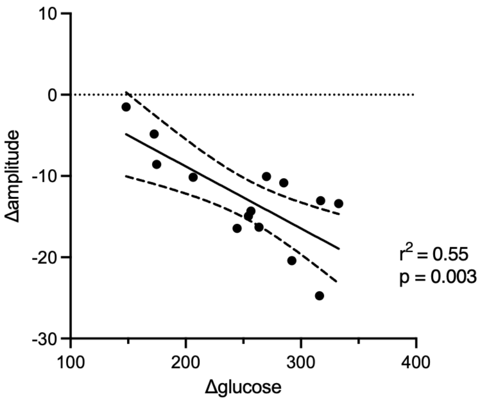


**B** (i)

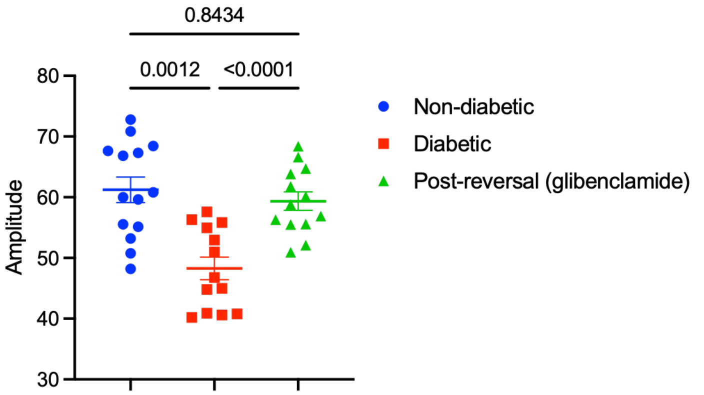


(ii)

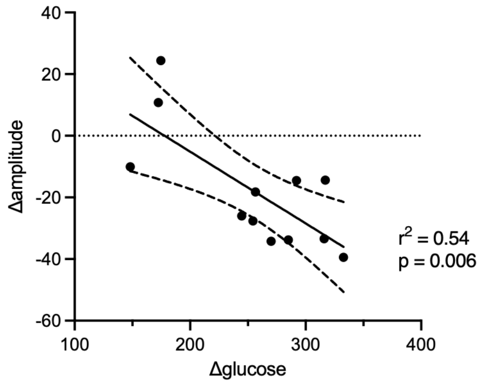

### Supplementary Figure 4

Representative activity profiles in LD (mean of 5 days activity, same individual in each time period). Blue dotted line shows outline of activity profile in non-diabetic mice for comparison to pre-induction baseline.

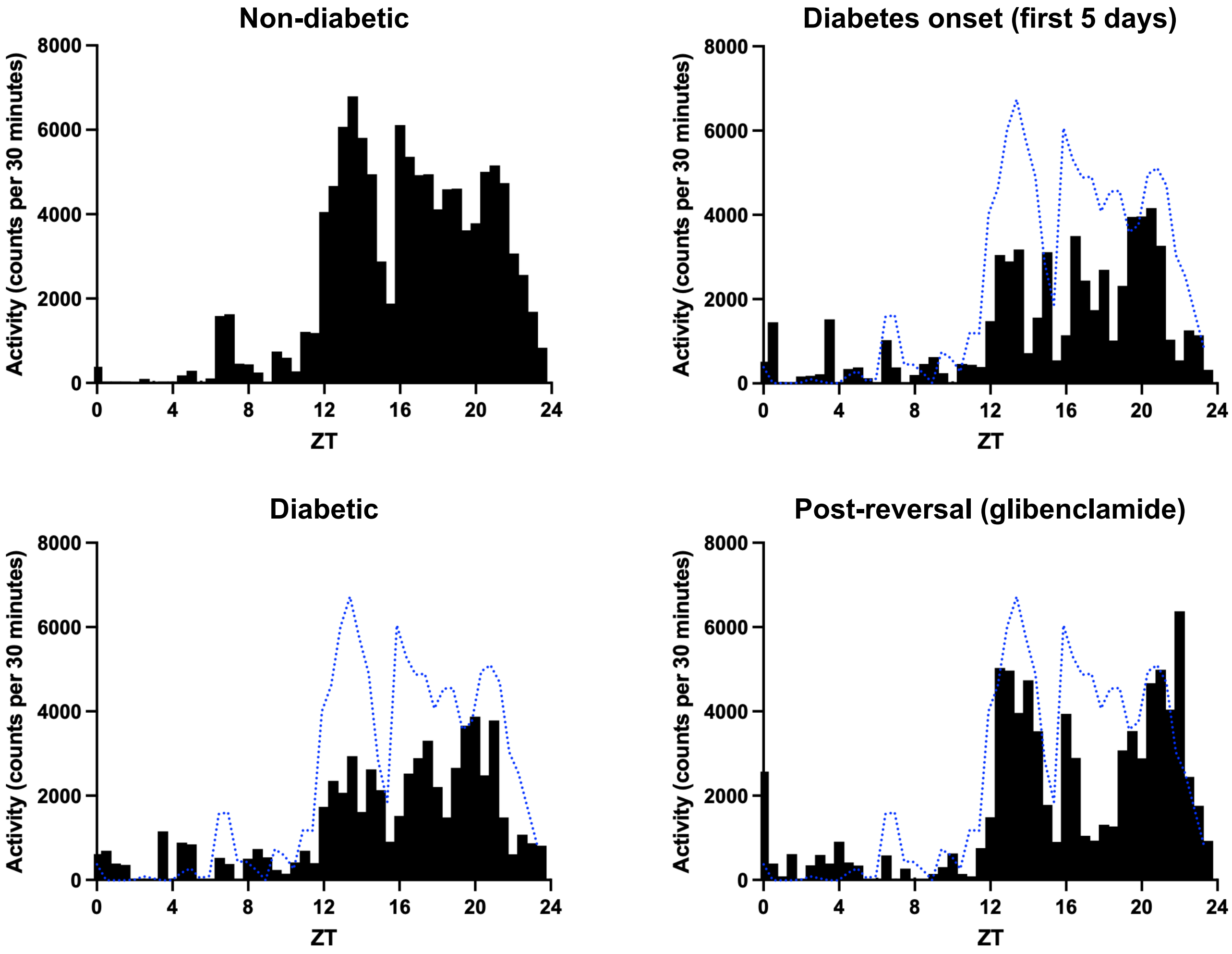

### Supplementary Figure 5

Amplitude of rest-activity rhythms in 6 C57BL/6 male mice before tamoxifen administration (days -3 to -1), immediately after injection with 50 mg/kg tamoxifen (days 0 to 2, where injection occurred on day 0), and following recovery from acute effects of the subcutaneous injection (days 3-5). Activity profile analysed across three consecutive 3-day windows.

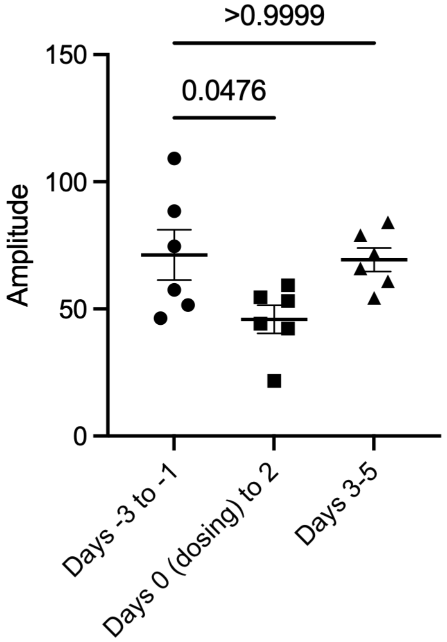


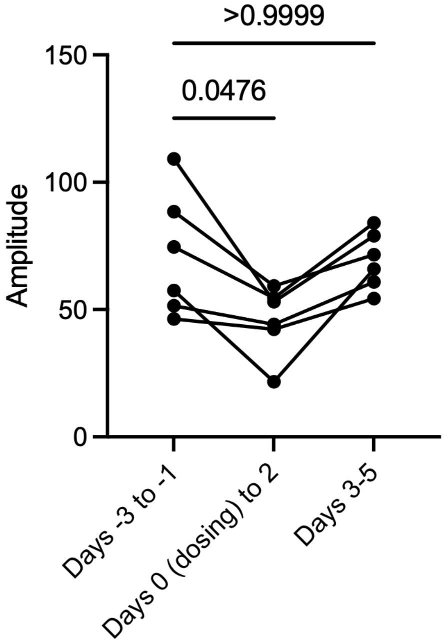

### Supplementary Figure 6

Activity analysis: intradaily variability (IV) and interdaily stability (IS) in (A) LD and (B) DD

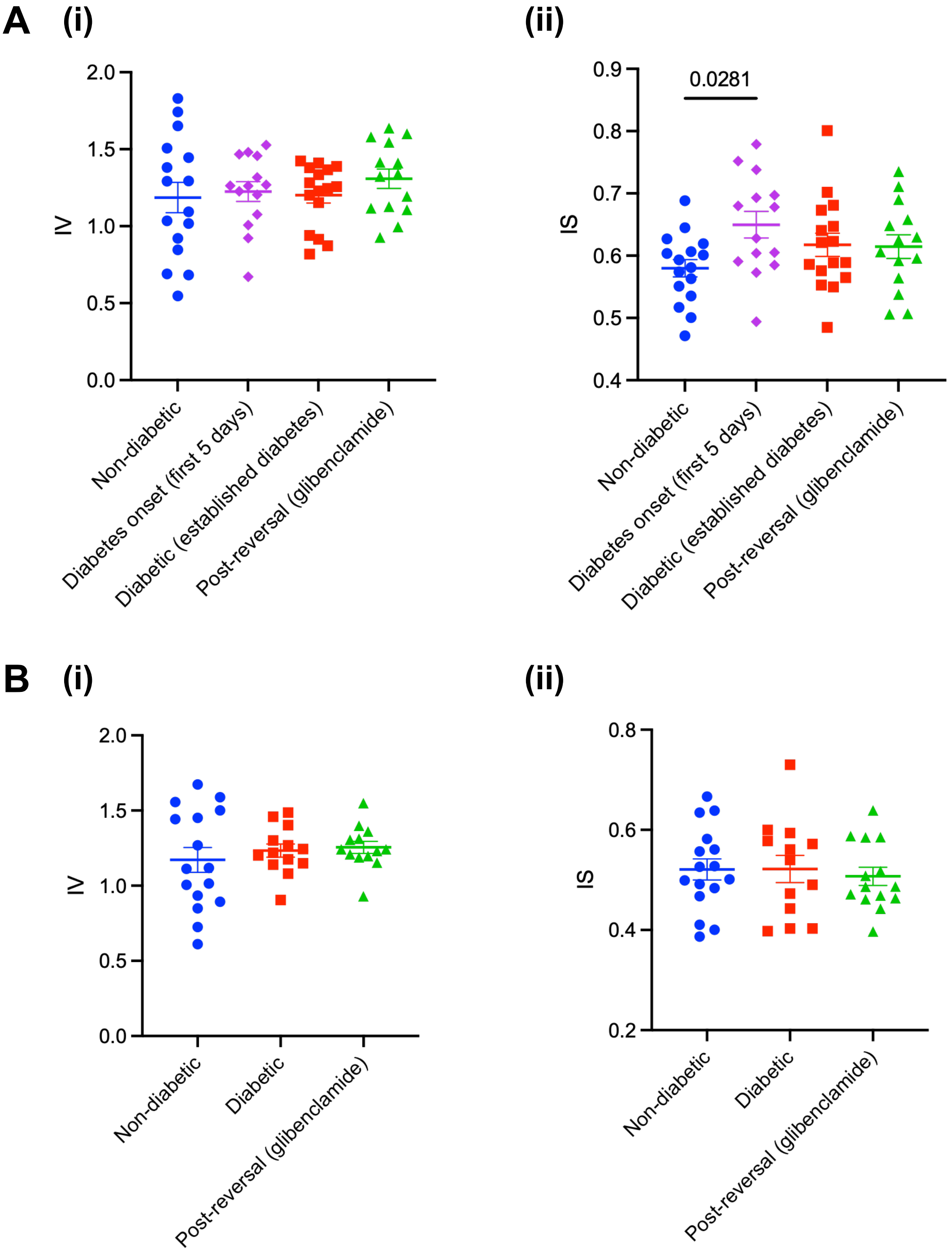

### Supplementary Figure 7

Representative actograms showing (A) DD, (B) 12/12 LD with 6h jet lag, and (C) DD with CT14 30min light pulse

#### A (i) Non-diabetic

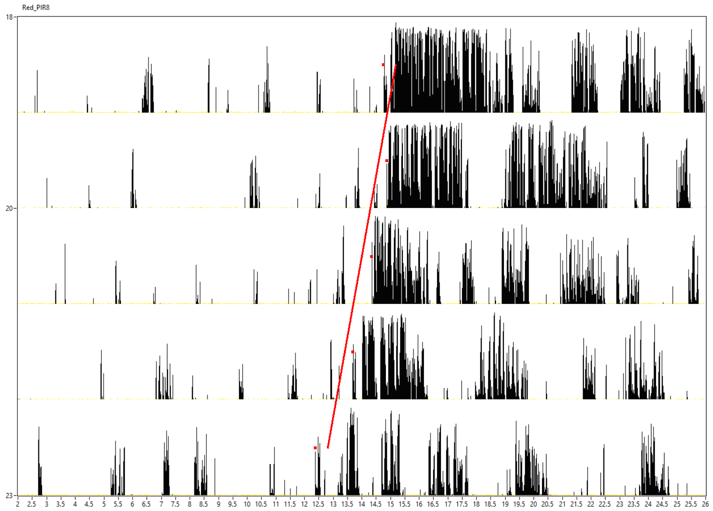


##### (ii) Diabetic

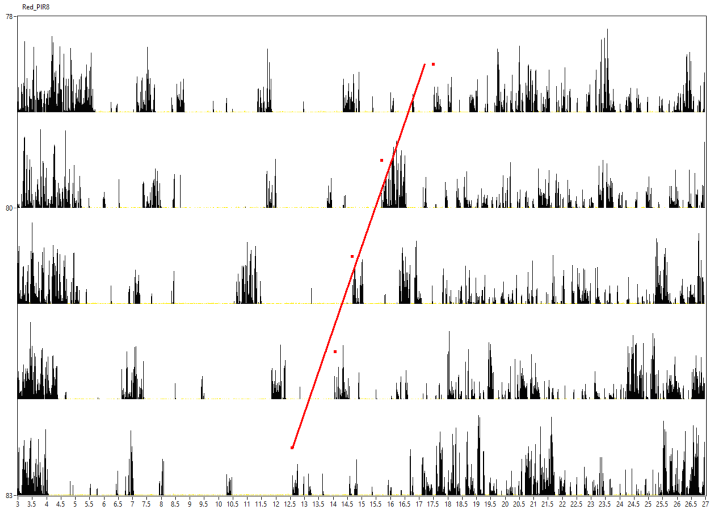


##### (iii) Post-reversal

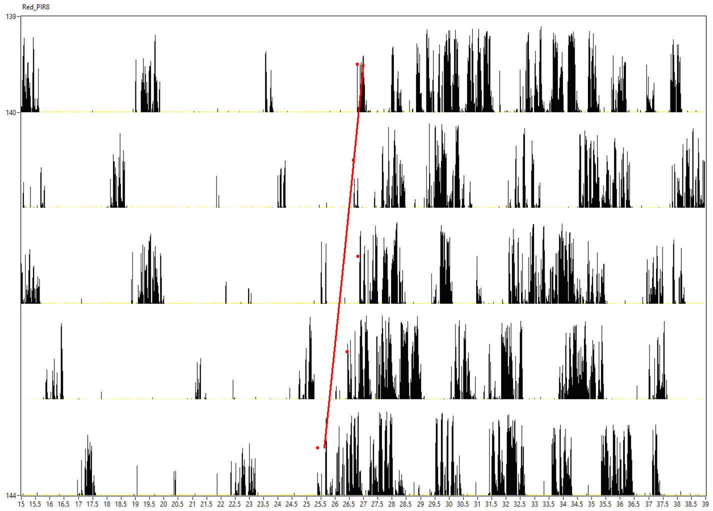

**B** **(i)** Non-diabetic

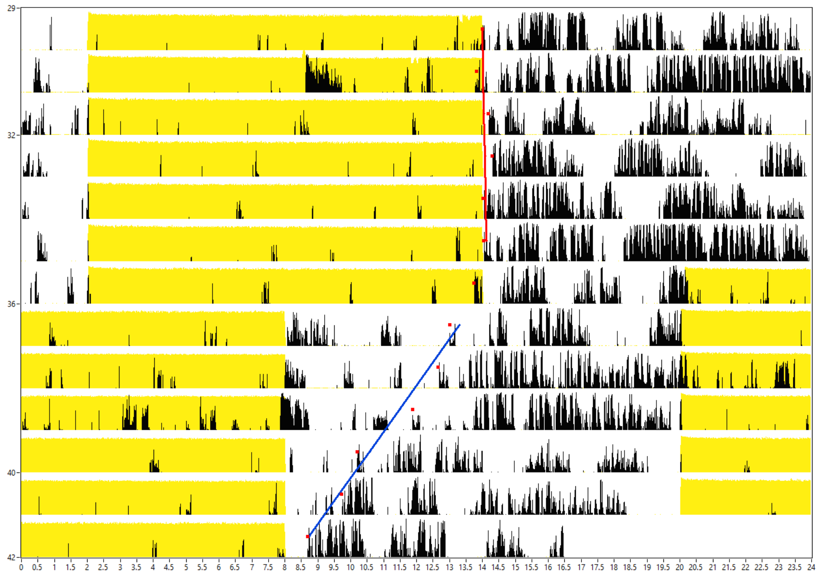


**(ii)** Diabetic

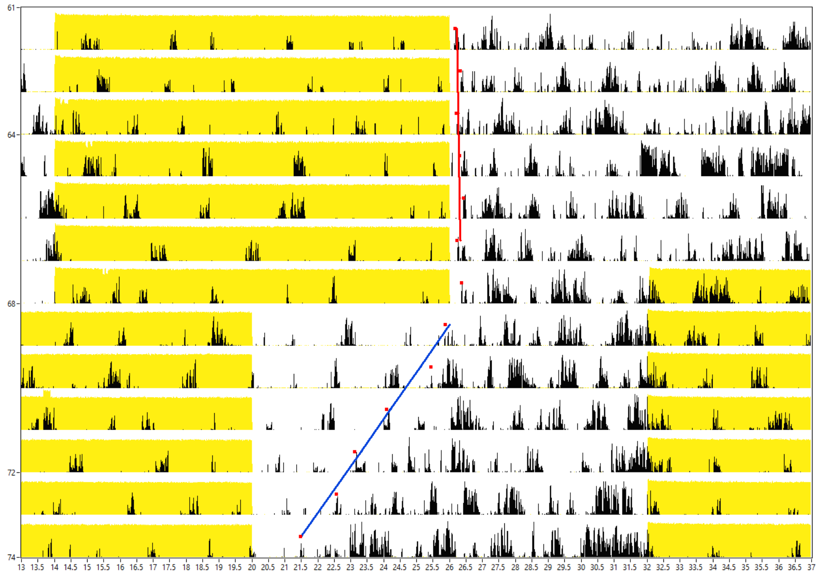


**(iii)** Post-reversal

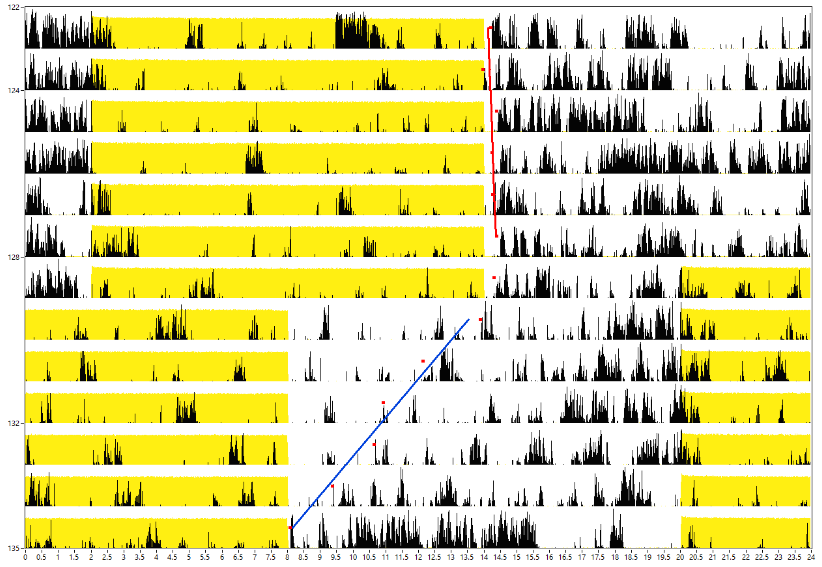

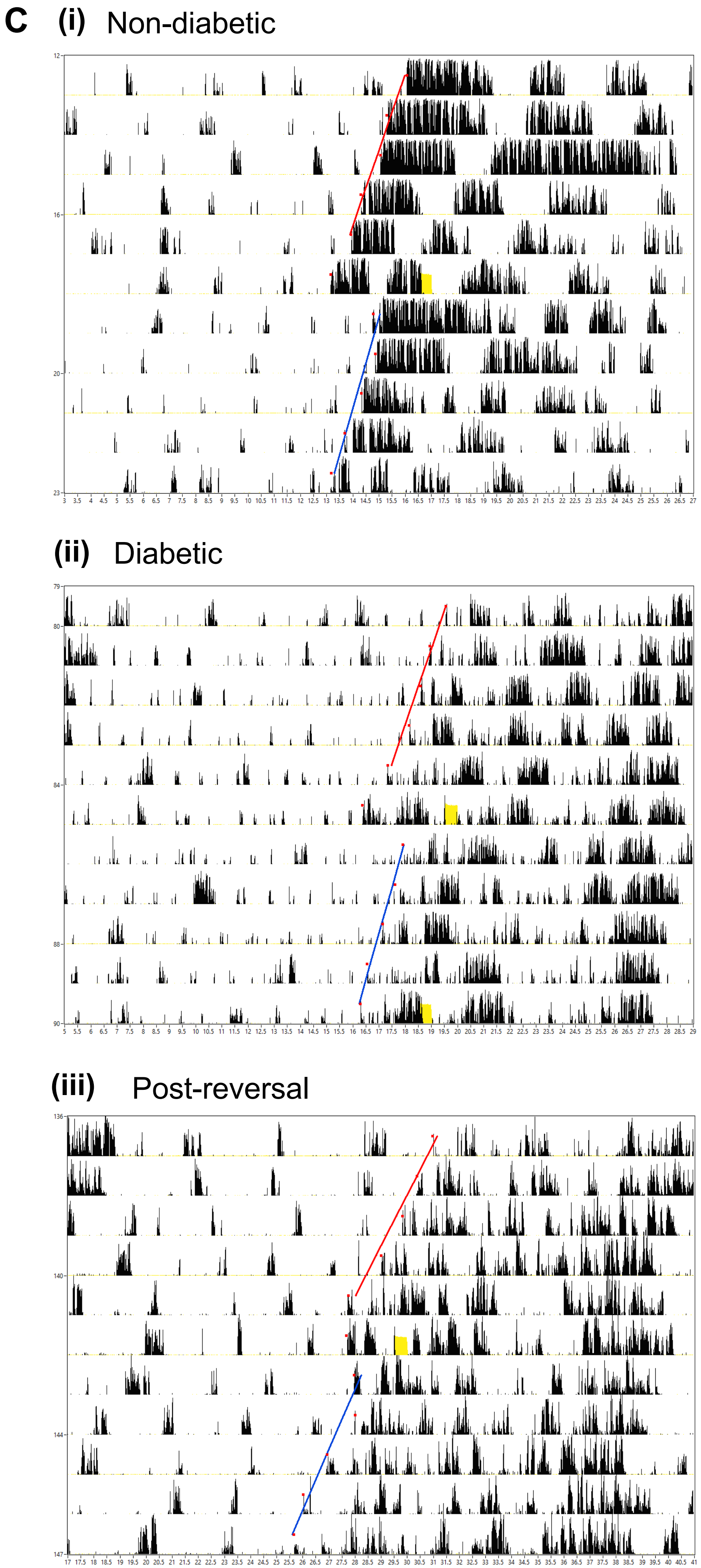
C
(i) Non-diabetic
(ii) Diabetic
(iii) Post-reversal

### Supplementary Figure 8

Scans of Western blot membranes

(A) Annotated representative blot showing molecular weight labels and each lane labelled with blood glucose concentration (mM) at time of death

(B) Membranes stained for O-GlcNAc

(C) Total protein stain corresponding to (B)

A

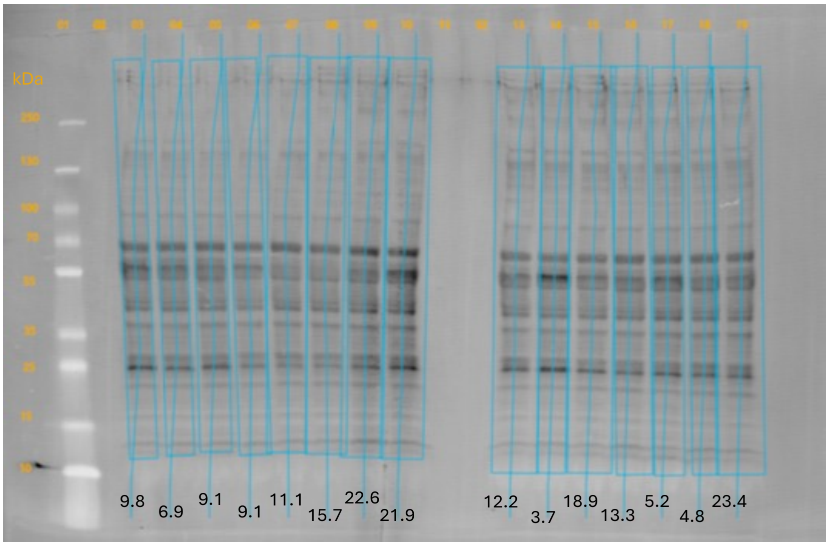


B

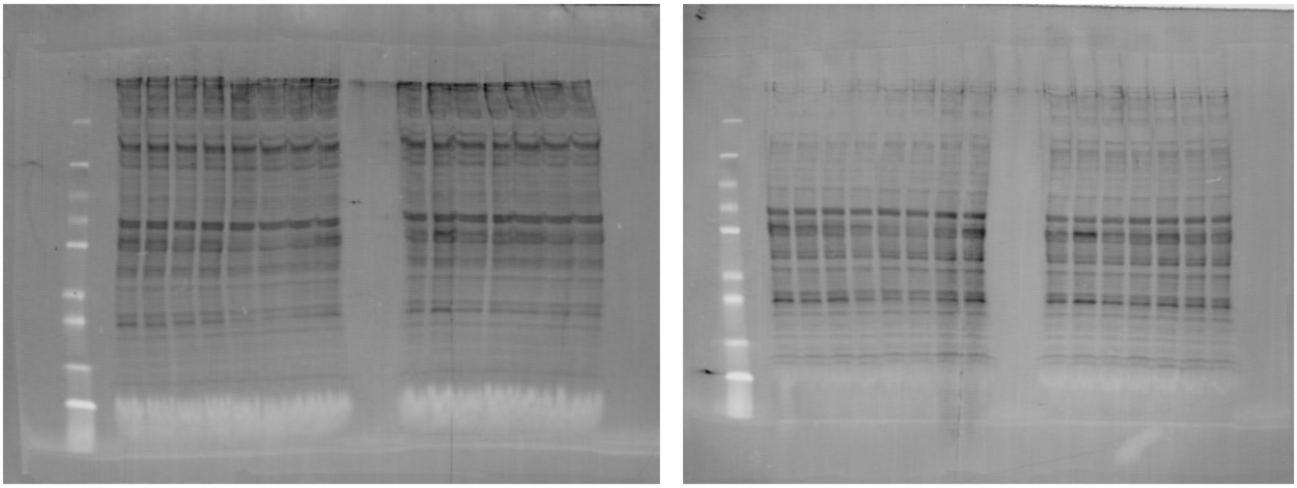

C

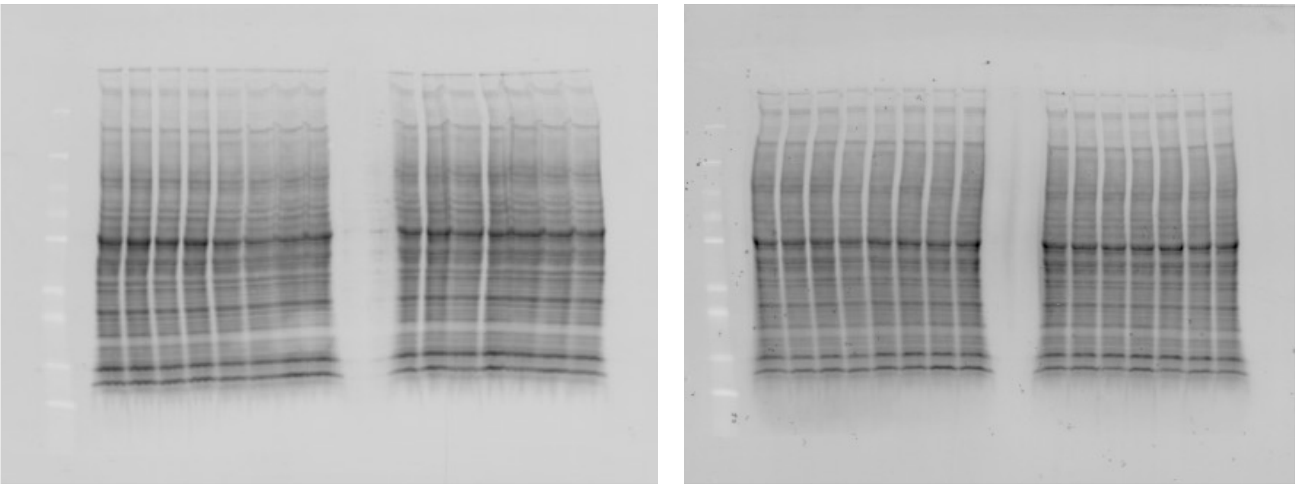

### Supplementary Figure 9

Brain glucose levels

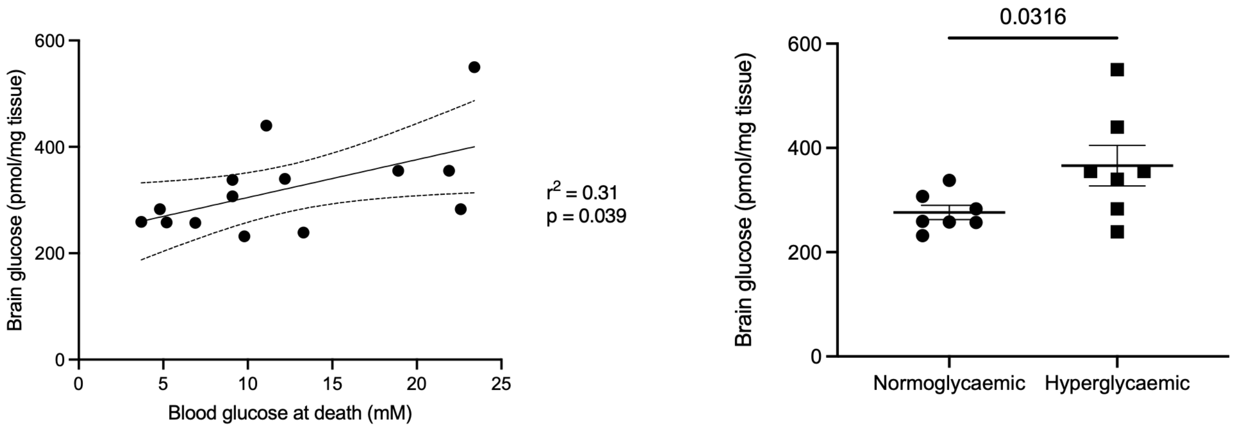
